# The Dutch Microbiome Project defines factors that shape the healthy gut microbiome

**DOI:** 10.1101/2020.11.27.401125

**Authors:** R. Gacesa, A. Kurilshikov, A. Vich Vila, T. Sinha, M.A.Y. Klaassen, L.A. Bolte, S. Andreu-Sánchez, L. Chen, V. Collij, S. Hu, J.A.M. Dekens, V.C. Lenters, J.R. Björk, J.C. Swarte, M.A. Swertz, B.H. Jansen, J. Gelderloos-Arends, Lifelines cohort study, M. Hofker, R.C.H. Vermeulen, S. Sanna, H.J.M. Harmsen, C. Wijmenga, J. Fu, A. Zhernakova, R.K. Weersma

## Abstract

The gut microbiome is associated with diverse diseases, but the universal signature of an (un)healthy microbiome remains elusive and there is a need to understand how genetics, exposome, lifestyle and diet shape the microbiome in health and disease. To fill this gap, we profiled bacterial composition, function, antibiotic resistance and virulence factors in the gut microbiomes of 8,208 Dutch individuals from a three-generational cohort comprising 2,756 families. We then correlated this to 241 host and environmental factors, including physical and mental health, medication use, diet, socioeconomic factors and childhood and current exposome. We identify that the microbiome is primarily shaped by environment and cohousing. Only ∼13% of taxa are heritable, which are enriched with highly prevalent and health-associated bacteria. By identifying 2,856 associations between microbiome and health, we find that seemingly unrelated diseases share a common signature that is independent of comorbidities. Furthermore, we identify 7,519 associations between microbiome features and diet, socioeconomics and early life and current exposome, of which numerous early-life and current factors are particularly linked to the microbiome. Overall, this study provides a comprehensive overview of gut microbiome and the underlying impact of heritability and exposures that will facilitate future development of microbiome-targeted therapies.

## Introduction

Alterations in gut microbiota composition and function are associated with a broad range of human health disorders, from gastrointestinal (GI) and metabolic diseases to mental disorders^1–3^. The influence of the gut bacteria and microbial pathways on host metabolism and immunity, together with the fact that the microbiota can be modified, have led to heightened interest in developing microbiota-targeted therapies^4,5^. In this context, however, the characteristics of a “healthy” microbiome remain largely unclear, as does the extent to which the gut microbiome is driven by intrinsic and external factors that might be amenable to microbiota-targeted therapies.

The capacity to define a “healthy” microbiome has been hampered by the large variation in microbiome composition between individuals and the large variability in the processing of faecal samples between studies. Population-based studies, including our previous study on ∼1500 Dutch volunteers^6^, have shown that this variation is partially accounted for by diet, lifestyle, host genetics and environmental factors, including early-life exposures^6–8^. However, while the number of microbiome studies has increased exponentially, in-depth integrative analyses of large standardized, well-phenotyped cohorts remain scarce, even though this integrative perspective is essential to disentangle meaningful host–microbiota associations and identify potential targets for microbiota-directed interventions.

To address these issues, we initiated the Dutch Microbiome Project (DMP) and analysed the gut microbiome composition and functionality of 8,208 individuals from the Lifelines study – a three-generational population cohort from a geographic region of approximately 11,400 km^2^ in the Northern Netherlands^9^. By exploring 241 host characteristics, we determined the impact of shared household, socioeconomic status, lifestyle and environmental exposures on the gut microbiota, which together explain 12.9% of the inter-individual variation in gut microbiota composition and 16.3% of the variation in microbiome functionality. Based on family relations, we estimated the heritability of the gut microbiota, which varies widely between different species (max H^2^ = 0.3). We also define the core microbiota and keystone species in the taxonomic networks of this Dutch population. After correcting for the impact of technical factors, age, sex, BMI and Bristol stool scale of faecal samples, we identify clear patterns of microbial taxa and pathway associations between multiple unrelated groups of diseases and with self-perceived health, which allows us to define features of a healthy or unhealthy microbiome and to predict the health status independently of comorbidities. Finally, we identified intrinsic and external factors, including environmental pressures (i.e. exposome), that influence the (un)healthy microbiome. Overall, this study provides a comprehensive large-scale exploration of the gut microbiome that will facilitate future investigations of microbiome-targeted therapies for improvement of long-term human health.

## Methods

### Population cohort and metadata collection

The Lifelines Dutch Microbiome Project (DMP) cohort has been developed as a part of the Lifelines cohort study. Lifelines is a multi-disciplinary prospective population-based cohort study using a unique three-generation design to examine the health and health-related behaviours of 167,729 people living in the North of the Netherlands. It employs a broad range of investigative procedures to assess the biomedical, socio-demographic, behavioural, physical and psychological factors that contribute to health and disease in the general population, with a special focus on multi-morbidity and complex genetics^9,10^. A total of 8,719 fresh frozen faecal and blood samples were collected from Lifelines participants in 2015 and 2016 to form the DMP cohort. Whole-genome shotgun sequencing was performed on 8,534 faecal samples, of which 8,208 were retained for downstream analysis after stringent quality control. Metadata information collected from the participants was grouped into the following categories: family structure, diseases, gastrointestinal (GI) complaints, general health score, medication use, anthropometrics, birth-related factors, reported childhood (< 16 years) exposures, current exposome (air pollutants, greenspace, urbanicity, pets and smoking), socioeconomic characteristics and diet (Supplementary table 2d).

### Informed consent

The Lifelines study was approved by the medical ethical committee from the University Medical Center Groningen (METc number: 2017/152). Additional written consent was signed by all DMP participants or their parents or legal representatives (for children aged under 18).

### Metadata

Metadata was collected by questionnaires and curated as described previously^10^ and below. We included 241 phenotypes from a broad range of categories, including socioeconomic factors, self-reported diseases and medications, quality of life, mental health, education and employment, nutrition, smoking, stress and childhood environmental factors. Questionnaires were developed and processed by the Lifelines cohort study^10^ as described at www.lifelines.nl. Additional in-depth data curation and acquisition was performed to assess dietary intake, air pollution and environmental exposures, medication use and gut health, as described below.

### Diet

Habitual diet was assessed through a semi-quantitative Food Frequency Questionnaire (FFQ) collected 4 years prior to DMP faecal sampling^10^. The FFQ was designed and validated by the division of Human Nutrition of Wageningen University using standardized methods^11^. It assesses how often a food was consumed over the previous month on a scale ranging from ‘never’ to ‘6-7 days per week’, along with the usual amount taken. The average daily nutrient intake was calculated using the Dutch Food Composition database (NEVO, RIVM) and a mono- and disaccharide-specific food composition table^12^, resulting in the generation of data on 21 dietary factors. Energy adjustment was performed by means of the nutrient density method^13^. Published dietary scores and inter-nutrient ratios were calculated as indicators of dietary quality and composition^14,15^.

To validate the assumed stability of FFQs across time^12,16^, we used questionnaires from 128,501 Lifelines participants to study diet consistency between the baseline questionnaire, collected 4 years prior to this study, and a second smaller-scope nutrient-specific questionnaire collected concurrently with DMP faecal sampling. Sixty-five dietary questions, reflecting consumption of major food categories such as fruits, vegetables, fish, meat, bread, grains and sweets, as well as special dietary regimes (e.g. vegan or macrobiotic diet), were compared between the first and second time point. The majority of dietary items were available for >44,000 individuals at both timepoints (Supplementary table 8). To quantify the consistency of answers and the degree of change in the 4-year period, we calculated the consistency (proportion of identical answers) and Euclidean distance between all participants who answered a question the same way at baseline and follow-up and the distance between participants who answered differently at follow-up (Supplementary table 8).

### Exposome

Elements of the exposome, neighbourhood urbanicity and income were assessed for the participant’s home address at the time of faecal sampling. Exposure to two air pollutants, particulate matter with aerodynamic diameter ≤ 2.5 μm (PM_2.5_) and nitrogen dioxide (NO_2_), was assigned based on land use regression models developed in the European Study of Cohorts for Air Pollution Effects (ESCAPE) project^17,18^. These estimates are based on measurement data from 2009 and reflect long-term ambient air pollution exposures^19^.

Greenspace was assigned using the Normalized Difference Vegetation Index (NDVI), which reflects the average density of green vegetation within a 100-meter circular buffer around the participant’s residential address. The NDVI was derived from a LANDSAT 5 (TM) satellite image taken in 2016 and captures the density of green vegetation at a spatial resolution of 30×30 m based on land surface reflectance of visible (red) and near-infrared parts of spectrum.

Neighbourhood urbanicity was assigned based on a five-category scale of surrounding address density developed by Statistics Netherlands (1 = very urban, ≥ 2500 addresses per km^2^ to 5 = very rural, < 500 addresses per km^2^, data for 2015)^20^. Neighbourhood income was considered a proxy of neighbourhood socioeconomic position and defined as the proportion of persons with low (< 40^th^ percentile) income (Statistics Netherlands, data for 2015)^20^.

### Stool characteristics, diseases and medication

Participants recorded a bowel movement diary, Bristol stool scale, daily medication use and GI symptoms daily for seven days in the week of stool sample collection, and these records were used to extract information on drug use, stool characteristics, stool frequency and GI symptoms during the week of stool collection. The validated ROME III questionnaires^21^ were used to characterize functional GI disorders, and participants were classified as having either no functional GI diseases or irritable bowel syndrome, functional diarrhoea, functional constipation or functional bloating. Information about the presence of other diseases was self-reported and collected using Lifelines questionnaires. Diseases were grouped into 11 disease categories. The presence of cancer was grouped into a separate category defined as “any cancer”, independent of cancer type. Non-alcoholic fatty liver disease fibrosis score^22^ and fatty liver index^23^ were calculated from the anthropometrics and blood measurements, as described previously^22,23^. Diseases with < 20 cases were excluded from further analysis. Self-reported medications were grouped into categories based on Anatomical Therapeutic Chemical classification (ATC codes) at the most-specific ATC level (5-digit ATC code if possible). ATC categories with < 20 users were grouped into a higher level (4-digit or 3-digit) ATC class, and categories with < 20 individuals that could not be grouped according to ATC classification were excluded from further analysis. In total, 62 drug groups were included (Supplementary table 2a).

### Sample collection, DNA extraction and sequencing

Faecal sample collection was performed by participants at home. Participants were asked to freeze stool samples within 15 min of stool production. The frozen samples were collected by Lifelines personnel, transported to the Lifelines biorepository on dry ice and stored at - 80°C until DNA extraction. Microbial DNA was isolated with the QIAamp Fast DNA Stool Mini Kit (Qiagen, Germany), according to the manufacturer’s instructions, using the QIAcube (Qiagen) automated sample preparation system. Metagenomic sequencing was performed at Novogene, China using the Illumina HiSeq 2000 platform to generate approximately 8 Gb of 150 bp paired-end reads per sample (mean 7.9 gb, st.dev 1.2 gb).

### Profiling microbiome composition and function

Metagenomes were profiled consistent with previous data analysis of 1000IBD^24^ and Lifelines-DEEP^10^ cohorts, as follows. KneadData tools (v0.5.1)^25^ were used to process metagenomic reads (in fastq format) by trimming the reads to PHRED quality 30 and removing Illumina adapters. Following trimming, the KneadData integrated Bowtie2 tool (v2.3.4.1)^26^ was used to remove reads that aligned to the human genome (GRCh37/hg19).

Taxonomic composition of metagenomes was profiled by MetaPhlAn2 tool (v2.7.2)^27^ using the MetaPhlAn database of marker genes mpa_v20_m200. Profiling of genes encoding microbial biochemical pathways was performed using the HUMAnN2 pipeline (v0.11.1)^28^ integrated with the DIAMOND alignment tool (v0.8.22)^28^, UniRef90 protein database (v0.1.1) and ChocoPhlAn pan-genome database (v0.1.1). As a final quality control step, samples with unrealistic microbiome composition (eukaryotic or viral abundance > 25% of total microbiome content or total read depth < 10 million) were excluded, leaving 8,208 samples for further analyses. Analyses were performed using locally installed tools and databases on CentOS (release 6.9) on the high-performance computing infrastructure available at our institution and using the MOLGENIS data platform^29^.

In total, we detected 1,253 taxa (4 kingdoms, 21 phyla, 35 classes, 62 orders, 128 families, 270 genera and 733 species) and 564 pathways in at least one of the samples in the quality controlled–dataset. To deal with sparse microbial data in the downstream analysis, we focused on bacterial and archaeal species/pathways with a mean relative abundance > 0.01% that are present in at least 5% of participants. This yielded 257 taxa (6 phyla, 11 classes, 15 orders, 30 families, 59 genera and 136 species) and 277 pathways. Together, these microbial features accounted for 97.86% and 87.82% of the average taxonomic and functional compositions, respectively.

Based on the abundance profiles of taxa that passed the filtering process, alpha diversity, as measured by richness and Shannon entropy, was calculated at family-, genus- and species-level using *specnumber* and the function *diversity*, respectively, in R package vegan (v.3.6.1). Rarefaction and extrapolation (R/E) sampling curves for estimation of total richness of species and genera in the population were constructed using a sample size–based interpolation/extrapolation algorithm implemented in the iNEXT package for R^30^.

### Profiling of bacterial virulence factors and antibiotic resistance genes

Metagenomes were searched for bacterial virulence factors (VFs) using the shortBRED toolkit (v0.9.5)^31^ and the database of Virulence Factors of Pathogenic Bacteria (VFDB) core dataset of DNA sequences (downloaded on 01/11/2018)^32^. The shortBRED tool shortbred_identify.py (v0.9.5) was used to identify unique markers for VFs, with the UniRef90 database (downloaded on 01/11/2018) used as negative control, and the shortbred_quantify.py tool (v0.9.5) was used to perform a quantification of these markers in metagenomes. Quantification of antibiotic resistance genes was performed using shortBRED tool shortbred_quantify.py (v0.9.5), with markers generated using shortbred_identify.py (v0.9.5) on the CARD database of bacterial antibiotic resistance genes (downloaded 01/11/2018)^33^, with the UniRef90 database used as negative control. This identified 190 VFs and 303 antibiotic resistance gene families (ARs), of which 47 VFs and 98 ARs were present in at least 5% of participants with relative abundance > 0.01%. These accounted for 95.22% of VF composition and 98.08% of AR composition, respectively.

### Estimation of heritability of microbiome

To accommodate covariate effects, we estimated the heritability of microbiome features using a variance components model implemented in the software POLY34,35. We first considered a base model in which variance is partitioned into a polygenic component, Vg, shared between individuals that is proportional to their kinship coefficient, and an environmental component, Ve, that is unique to each individual. Thus, if Y is the measured trait, the variance of Y is Var(Y) =Vg+Ve, and its broad heritability is H2=Vg/(Vg+Ve). Of note, this measure reflects the overall impact of genes on the phenotype, thus including all potential models of action, as opposed to narrow heritability, which only includes additive effects. After fitting this base model, we also considered two refined models that included i) an additional variance component to model a shared environment for people belonging to the same family, Vh, and ii) a covariate with values 0/1 that distinguishes family members currently living in the same house from those who do not. We compared the significance of these two models to the basic model using a likelihood ratio test and found the base model to be the most appropriate fit for all microbiome traits.

We restricted the analysis of heritability to the relative abundances of the 242 microbial taxa present in at least 250 individuals and focused on 4,745 individuals in 2,756 families in which at least two individuals had available microbiome data. In total, the analysis included 2,756 parent-child pairs, 530 sibling pairs and 815 pairs with second-degree or more distant relationships. All models were adjusted for age, sex, BMI, read depth and stool frequency, and values of relative abundances of taxa were transformed using the centred log-ratio (clr) transformation^36^. Benjamini-Hochberg correction was used to control the multiple testing false discovery rate (FDR), and results with FDR < 0.1 were considered significant.

### Estimation of the effect of co-housing on microbiome

We estimated the effect of co-housing on microbiome composition, function, ARs and VFs by comparing beta-diversities of the microbiomes of co-housing study participants (1,710 unrelated pairs, 285 parent–child pairs and 144 sibling pairs) to those of participants who did not share housing (2,000 unrelated pairs, 301 parent–child pairs and 299 sibling pairs). Microbiome distance was calculated for every pair using Bray-Curtis dissimilarity, and mean dissimilarities within groups were compared using Mann-Whitney U tests using the Benjamini-Hochberg correction to control multiple testing FDR. Results were considered significant at FDR < 0.05.

### Calculation of microbiome–phenotype associations

The microbiome composition variance explained by phenotypes was calculated by permutational multivariate analysis of variance using distance matrices, implemented in the *adonis* function for R package *vegan* (v.2.4-6), using 20,000 permutations and a Bray-Curtis distance matrix calculated using relative abundances of microbial species. A separate analysis was performed to calculate the microbiome functional potential explained by phenotypes using equivalent methodology. The functional dissimilarity matrix was calculated using the Bray-Curtis dissimilarity index calculated on the relative abundances of MetaCyc microbial biochemical pathways.

Prior to association analysis of phenotypes and microbiome features, the microbiome data was transformed using the clr transformation. The geometric mean for clr transformation of relative abundances of taxa was calculated on species-level and applied to higher levels. The associations between phenotypes and microbial features (microbial taxa, MetaCyc functional pathways, CARD and VFDB entities) were calculated using linear regression, adjusting for age, sex and BMI of the individual along with Bristol stool scale of the faecal sample and technical factors (DNA concentration, sequencing read depth, sequencing batch and sampling season). Benjamini-Hochberg correction was used to control for multiple testing with the number of tests equal to the number of tested feature–phenotype pairs. Results were considered significant at FDR < 0.05.

### Quantification of microbiome–disease–drug interactions

To disentangle interactions between the gut microbiome, medication and diseases, we explored the effect of a selection of drugs and the common diseases for which these drugs are used: functional GI disorders and proton pump inhibitors, type 2 diabetes and antidiabetic drugs, and depression and selective serotonin reuptake inhibitors. We extracted subsets of participants with the disease and those who report using disease-associated medication and used these subsets to construct multivariate models including both medication use and disease, additionally adjusting for other factors as described above in *Calculation of microbiome-phenotype associations*.

### Definition of core microbiome and prediction of keystone microbiome features

To identify core microbial species and pathways, we used a bootstrapping-based selection approach. We randomly sampled 1% to 100% of the samples of the cohort a hundred times and calculated the standard deviation of the presence rate of each microbial species/pathway at different sampling percentages. Microbial features with a presence rate of more than 95% of samples were defined as the core microbiome, and we identified 9 core microbial species and 143 core microbial pathways (Supplementary table 1a,b).

To analyse microbiome community structure, we constructed microbial species and pathway co-abundance networks using SparCC, as previously published^37,38^. Relative abundances of taxa were converted to estimated read counts by multiplying abundance percentages by total sequenced reads per sample after quality control. For pathway analysis, the read counts (RPKM) from HUMAnN2 were directly used for SparCC. Significant co-abundance was controlled at FDR 0.05 level using 100 permutations. In each permutation, the abundance of each microbial feature was randomly shuffled across samples. In this way, we obtained 6,473 species and 55,407 pathway co-abundances at FDR

< 0.05. Features that ranked in the top 20% in the number of network connections (node degree) were considered keystone species or pathways, resulting in 28 keystone species and 53 keystone pathways (Supplementary table 1e).

### Identification of microbiome clusters

To identify microbial clusters and assess the presence of gut enterotypes in our cohort, we performed the partitioning around the medoid method on the relative abundances of microbial species and used the Calinski-Harabasz index to select the optimal number of clusters, as previously published in a study of gut enterotypes^39^. Enrichment of phenotypes in each cluster was assessed by logistic regression in R.

### Calculation of microbiome signatures predictive of diseases and health

We calculated the microbial signatures predictive of the 36 most common (N_cases_>100) diseases in our dataset. In addition, we defined a “healthy” phenotype as an absence of any self-reported disease. Using this definition, 2,937 (36%) out of 8,208 individuals were defined as “healthy”.

To build prediction models for common diseases, the dataset was randomly split into training (90%) and test (10%) sets. Next, we performed elastic net L1/L2 regularized regression (R package *glmnet* v.4.0) on the training set, using Shannon diversity, clr-transformed microbial taxa, clr-transformed MetaCyc bacterial pathways and age, sex and BMI as fixed covariates (not penalized in the models). The model for each disease was calculated independently using five-fold cross-validation to select the optimal lambda penalization factor (at L1/L2 mixing parameter alpha fixed at 0.5). The lambda with minimal cross-validation error was used in the downstream analysis.

In total, we defined three probabilistic models: a “null” signature that only includes effects of general covariates (age, sex and BMI), a “microbiome” signature that includes all selected microbiome features and a “combined” signature that includes both the effects of microbiome features and general covariates.

### Data availability

Raw sequencing data and corresponding metadata that support the findings of this study are available on request from the corresponding author R.K.W. The participant metadata are not publicly available as they contain information that could compromise research participant privacy/consent.

The authors declare that all other data supporting the findings of this study are available within the paper and its supplementary information files.

## Results

### Overall picture of gut microbial composition

In this study we characterized the composition and function of the gut microbiota of 8,208 individuals from the Northern Netherlands. Participants were of a wide age range (8-84 years), 57.4% were female and 4,745 individuals clustered into 2,756 families. The participants were largely (99.5%) of Dutch European ancestry (Fig.1a, Supplementary table 2d).

A total of 1,253 taxa (4 kingdoms, 21 phyla, 35 classes, 62 orders, 128 families, 270 genera and 733 species) and 564 metabolic pathways were present in the dataset, of which 257 taxa and 277 pathways had a relative abundance higher than 0.01 and were present in > 5% of individuals. Our sample size allowed us to capture ∼100% of the estimated microbial functional potential (including biochemical pathways, virulence factors and antibiotic resistance genes) and microbial taxa at level of genus or higher. By subsampling the cohort, we estimated that the presence rates of these microbial features become stable when 40% or more of the cohort is sampled (∼3,300 samples). However, we also observed that the number of microbial species discovered continued to increase with increasing sample size, with the total number of species in the population estimated to be 612 (standard deviation = 20) at 25,000 samples, suggesting that the sampled population contains rare undiscovered microbial species (Fig. 1b, Supplementary fig. 1). Gut microbiota composition was found to be highly variable across the population, with the relative abundance of the phylum Bacteroidetes, for example, ranging from 5% to > 95% (Supplementary fig. 2c). However, the most abundant microbial pathways were largely stable across the population (Supplementary fig. 2d).

**Figure 1:**
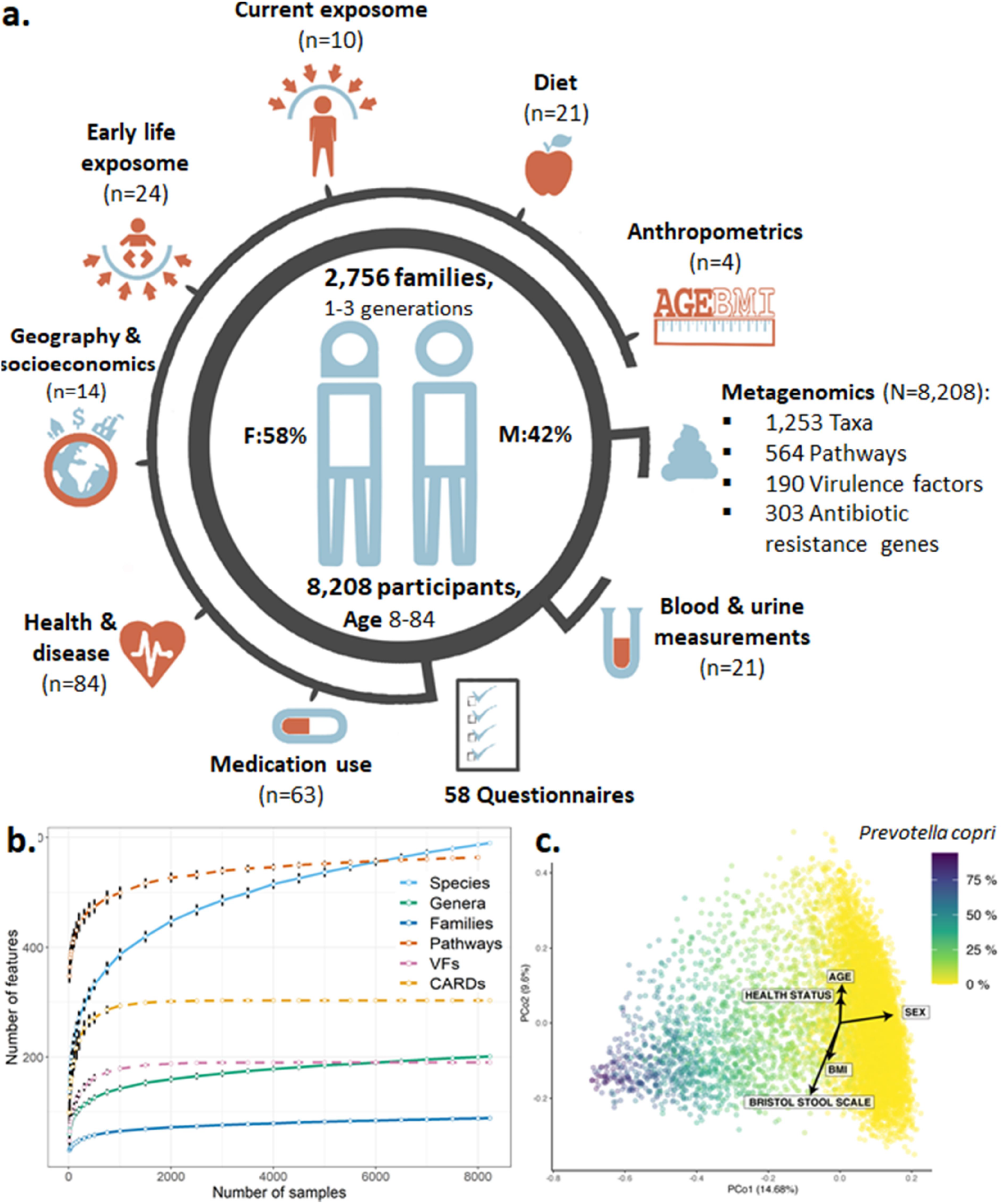
Summary of the Dutch Microbiome Project. **a**, Graphical summary of the cohort and overview of available metadata (n = number of variables collected, N = sample size). **b**, Number of microbial features discovered in relation to sample size. Error bars denote the standard deviation of 100 resamplings. **c**, Biplot of principal coordinate analysis visualising the beta-diversity of the microbiome data. Colour indicates relative abundance of Prevotella copri. Arrows indicate the influence of self-reported health, anthropometrics and faecal sample metadata. VFs: bacterial virulence factors, CARD: antibiotic resistance gene families

### Core microbes and pathobionts are keystone species in the gut ecosystem

To pinpoint bacterial species and pathways potentially critical for the organisation and maintenance of the gut ecosystem, we investigated 8,208 samples in our cohort for bacteria present in > 95% of individuals (*core microbes*) and for bacteria that form the central nodes in bacterial co-abundance networks *(keystone features*)^38,40^. We identified 9 core species (*Subdoligranulum sp*., *Alistipes onderdonkii, A. putredinis, A. shahii, Bacteroides uniformis, B. vulgatus, Eubacterium rectale, Faecalibacterium prausnitzii* and *Oscillibacter sp*.) that are highly consistent with those found in previous studies of UK, US and European and non-western populations (Supplementary Table 1a)^6,41–44^. We also identified 28 species and 53 pathways as potential keystone features defined by more than 109 and 337 significant co-abundances, respectively (false discovery rate (FDR) < 0.05).

We observed that 5 of the 9 core microbial species (*A. putredinis, A. shahii, F. prausnitzii, Oscillibacter sp*. and *Subdoligranulum sp*.) are also keystone species, implying that these 5 microbes are not only highly prevalent, they play central roles in the gut microbiome ecosystem in the Dutch population. For example, *F. prausnitzii*, a major butyrate producer that is depleted in many chronic diseases^45,46^, had a significant co-abundance with the majority of *Bacteroidetes* and *Bifidobacterium* species (Supplementary table 1e). In addition to these core species, we identified potential keystone species with low prevalence in the population (prevalence ≤ 0.1), including *Ruminococcus gnavus* and multiple species from genus *Clostridium*. These keystone bacteria were further observed to be positively associated with multiple diseases in the current study (Supplementary table 3b), as also seen in previously published studies^47,48^.

### Prevotella copri defines microbiome clusters

To evaluate if gut microbiomes in our cohort show distinct clusters, we examined the data using principal coordinate analysis (PCoA) and identified that the first principal coordinate is largely driven by *Prevotella copri* (rSpearman=0.68, P=3.6×10^−180^, Fig. 1c, Supplementary table 9). This bacterium is bimodally distributed in our cohort and defines two robust clusters based on its presence or absence (Supplementary fig. 3a,b). It was previously suggested that a gut microbiota with a low abundance of *Prevotella* is associated with higher incidence of irritable bowel syndrome (IBS)^49^, and we also observe that the cluster with a high abundance of *P. copri* has a lower risk of IBS (OR = 0.72, 95% CI 0.86-0.61) and is positively associated with general health (OR = 1.24, 95% CI 1.40-1.11, FDR < 0.05, Supplementary fig. 3c). While previous studies have reported distinct enterotypes dominated by *Bacteroides, Prevotella* and *Ruminococcaceae*^39,50^, we only observed two such clusters in our cohort, possibly because our cohort is ethnically uniform and comes from a constrained geographic area.

Unlike microbiome composition, the PCoA of functional potential was not dominated by any single pathway, and the top features explaining variance were found to be queuosine biosynthesis, peptidoglycan biosynthesis and L-isoleucine biosynthesis pathways (Supplementary table 9).

### Microbiome is largely determined by cohabitation, but core species are heritable

We next explored the relative contributions of family structure, co-housing and other exposome factors in shaping the gut microbiome. We utilised the multi-generational family structure of our cohort to estimate the heritability of microbial taxa and identified 31 heritable taxa (12.8% of the tested taxa) at FDR < 0.1 (Fig. 2a,b, Supplementary table 5). The highest heritable values were observed for *Sutterella wadsworthensis* (H^2^ = 0.3) and multiple species from genera *Bacteroides, Collinsella* and *Phascolarctobacterium* (H^2^ ∼0.21). While comprising a relatively minor part of the microbiome, the heritable taxa were found to be enriched within the set of core microbes (3/9 core taxa were heritable, Chi-squared test p-value < 1.0e-6) and to have significantly higher prevalence and abundance in the population compared to non-heritable taxa (Mann-Whitney U test p-values < 1.0e-6). In addition, 11/31 heritable taxa were positively associated with health, whereas only 3/31 heritable taxa were negatively associated with health (Supplementary table 5).

We further investigated the effect of cohabitation versus heritability by comparing the microbiome composition of first-degree relatives (parent–child pairs and siblings) living together to those of related pairs living separately and unrelated cohabiting participants (cohabiting partners) and to those of people living separately (randomly selected unrelated participants). The microbiomes of cohabiting pairs were more similar than the microbiomes of participants living separately, regardless of the relatedness of these pairs, with parent– child pairs, sibling pairs and unrelated partners all having microbiomes significantly closer to each other than randomly sampled non-cohabiting pairs (Fig. 2c, p-values < 1.0e-5). Microbiomes of cohabiting parent–child and sibling pairs were significantly more similar than equivalent pairs living separately, while related pairs living separately were not significantly different from random unrelated pairs (Fig. 2d,e). We further observed similar patterns in the compositions of microbial pathways, virulence factors and antibiotic resistance genes (Supplementary fig. 4a-c).

**Figure 2:**
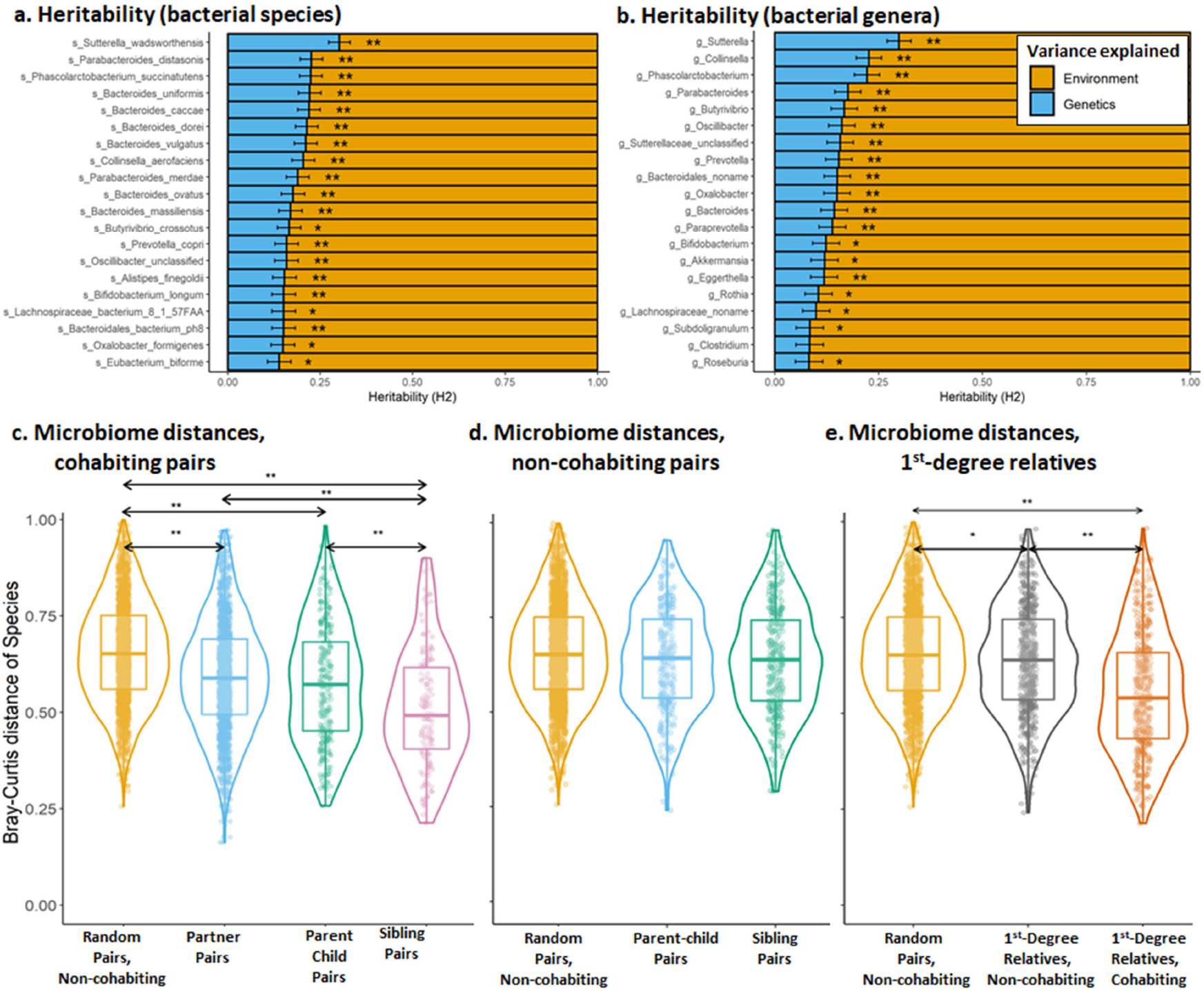
Heritability and impact of cohabitation on the gut microbiome. **a**, Top heritable species. **b**, Top heritable genera. ** taxa significantly heritable at FDR < 0.1. * taxa with nominally significant heritability (p-value < 0.05). Panels **c, d** and **e** show pairwise microbiome distance comparisons. Bray-Curtis dissimilarities calculated using microbial species of groups of **c**) random, non-cohabiting pairs compared to cohabiting partners, parent-child pairs and sibling pairs, **d**) random pairs compared to non-cohabiting parent-child and sibling pairs and **e**) random pairs compared to non-cohabiting first-degree relatives and cohabiting first-degree relatives. Significantly different groups: ** for FDR < 1.0e-5 and * for FDR < 0.05.

These results indicate that whole-microbiome composition is significantly influenced by cohabitation, with genetics playing a less important role. However, a minor proportion of the gut microbiome, e.g. species from genera *Sutterella* and *Collinsella*, is significantly heritable.

### Overview of microbiome–phenotype associations

We then explored the associations of microbial features to 241 individual measures including technical factors, anthropometrics, early-life and current exposome, diet, self-reported diseases and medication use, medical measurements and socioeconomics (Fig. 3a). These phenotypes explained 12.9% of microbiome taxonomic composition and 16.3% of microbiome function, with the largest contribution coming from technical factors, stool characteristics, diseases, medication use and anthropometrics (Fig. 3c, Supplementary tables 4a-c).

**Figure 3:**
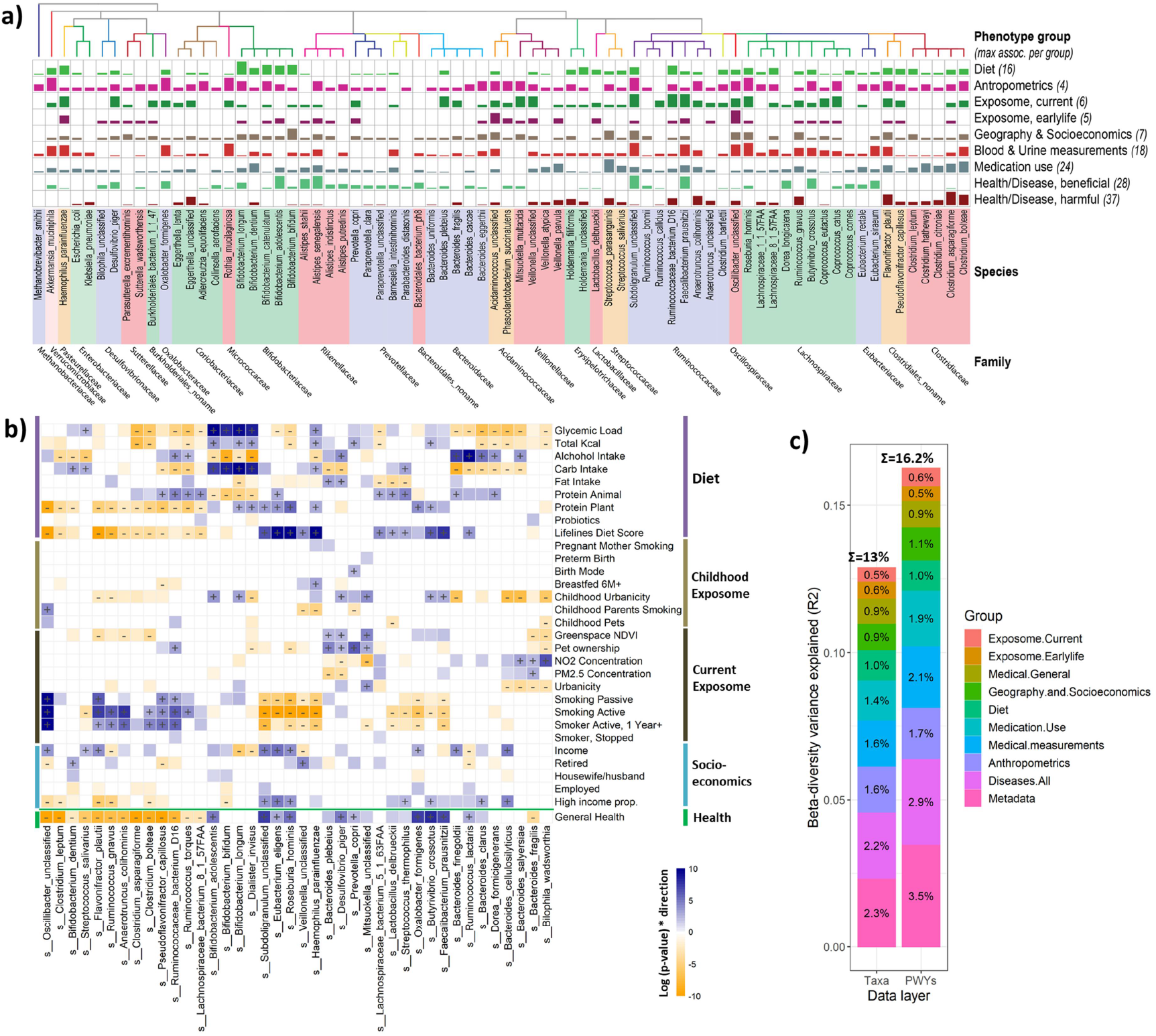
Microbiome–phenotype associations. **a**, Selected study-wide-significant associations (FDR < 0.05) per phenotype group, clustered by taxonomy. Bar height indicates the number of associations relative to the maximal number of associations for the phenotype group. **b**, Microbiome–phenotype associations for diet, childhood and current exposome and socioeconomics in comparison to healthy microbiome signature. Microbial species are clustered by association p-value using hierarchical clustering and coloured by direction of association. Study-wide significant associations (FDR < 0.05) are marked with + or -. Coloured associations without a mark indicate nominally significant associations (p-value < 0.05). **c**, The variance in microbiome composition and function explained by phenotype groups.

After correcting for technical factors, we observed 4,530 associations of phenotypes with taxa, 5,224 with pathways, 1,848 with antibiotic resistance genes and 385 with virulence factors (Supplementary table 3g, Supplementary fig. 5). Individually, the largest number of associations were observed for keystone and core taxa, including *Flavonifractor plautii, F. prausnitzii, Alistipes senegalensis* and species of genera *Clostridium* and *Subdogranulum* (Supplementary tables 3b,h). Below, we describe a selection of association results. Supplementary tables 3a-h provide a complete overview.

Age, sex and BMI ranked among the top phenotypes in our analysis of interindividual variation of beta-diversity, explaining 0.6%, 0.53% and 0.32%, respectively, as did individual microbiome feature associations (Supplementary tables 3a-e). Bristol stool scale explained the largest proportion of beta diversity for any single factor in our cohort (R^2^ = 0.77%, FDR = 0.012), and the sampling season also explained significant proportion of variance (R^2^ = 0.36%, FDR = 0.012, Supplementary table 4a), both highlighting the importance of assessing faecal sample consistency and collection time-frame effects in microbiome studies. Based on these results, we included corrections for age, sex, BMI, Bristol stool scale and sampling season into all our association models alongside the corrections for technical factors described above (Supplementary tables 3a-e).

### Definition of healthy and disease-associated gut microbiome

To define the microbiome signatures of health and disease, we associated microbiome features to self-reported health and 81 diseases that had at least 20 cases in the cohort, including GI and hepatological, cardiovascular and metabolic, mental and neurological, pulmonary, and other disorders. We identified 1,206 significant health/disease associations with bacterial taxa, 1,182 with microbial pathways, 390 with antibiotic resistance gene families and 76 with bacterial virulence factors (FDR < 0.05, Supplementary tables 3b-f). Different diseases showed varying numbers of associated microbes, with the strongest signatures observed for cardiovascular and metabolic disorders, such as non-alcoholic fatty liver disease and Type 2 Diabetes (T2D), and for GI disorders including Inflammatory Bowel Disease (IBD) and IBS (Supplementary table 3f). We observed consistent microbiome– disease patterns across the majority of diseases (Fig. 4), allowing us to pinpoint microbiome signatures shared between different/independent diseases as well as features that define a healthy (i.e. *absence of disease*) microbiome.

The shared microbiome signatures of disease (described in further detail in the **Supplementary Discussion**) mainly consisted of increases in species from *Anaerotruncus, Ruminococcus, Bacteroides, Holdemania, Flavonifractor, Eggerthella* and *Clostridium* genera and decreases in *Faecalibacterium, Bifidobacterium, Butyrivibrio, Subdoligranulum, Oxalobacter, Eubacterium* and *Roseburia*. Gut microbiome pathways shared across diseases with different aetiologies mainly consisted of increases in biosynthesis of L-ornithine, ubiquinol and menaquinol, enterobacterial common antigen, Kdo-2-lipid-A and molybdenum cofactor and decreases in biosynthesis of amino acids, deoxyribonucleosides and nucleotides, anaerobic energy metabolism and fermentation to short chain fatty acids (mainly butanoate). Furthermore, virulence factors were increased in some diseases, including T2D and GI disorders, with the largest effect observed for bacterial adherence and iron-uptake factors (VF036, VF0228, VF0236, VF0404 and VF0394).

**Figure 4:**
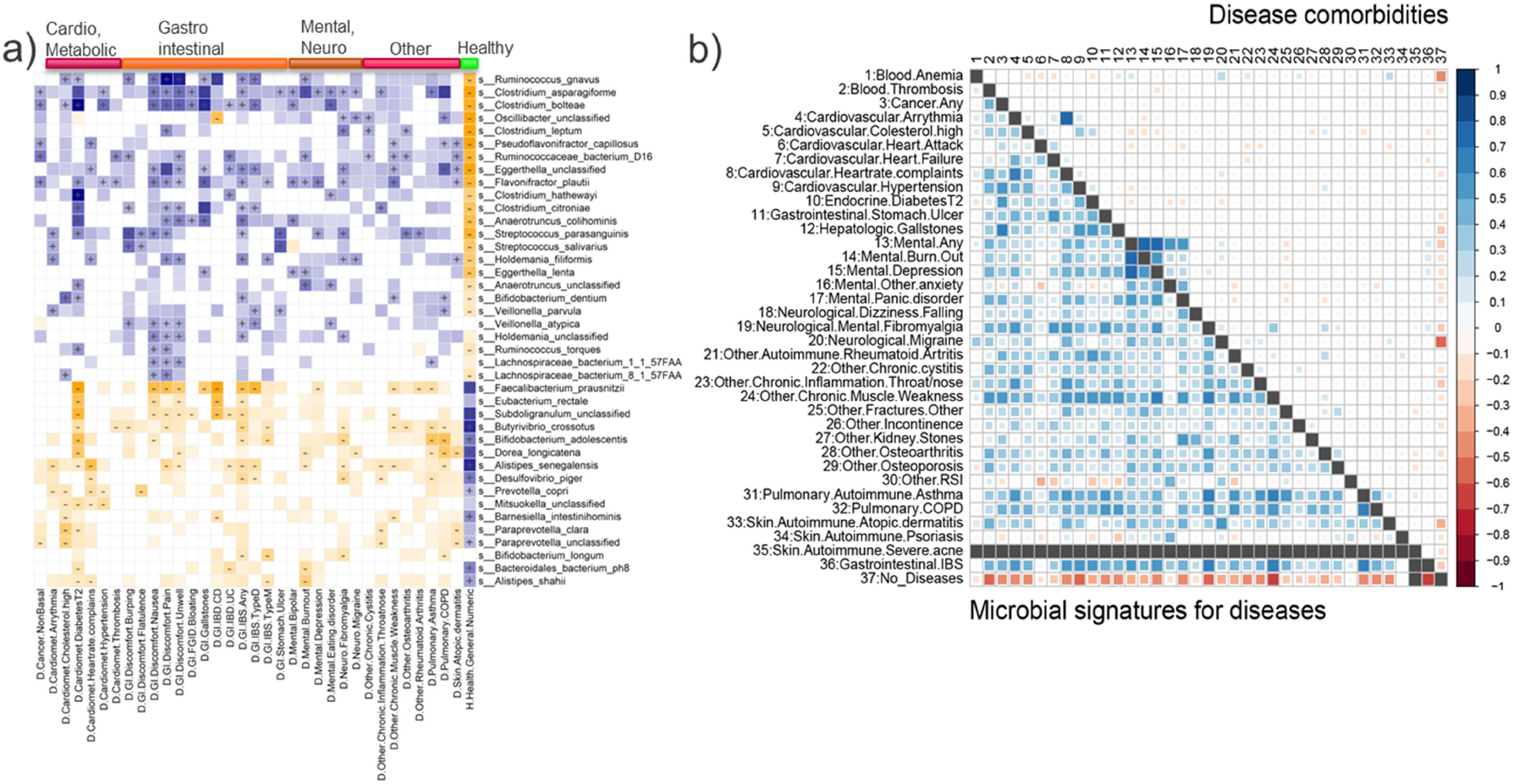
Microbiome signatures of health and diseases. **a**, Heatmap of microbial species associated with categories of diseases and health status. Diseases are sorted and labelled by disease type. Microbial species are clustered by association p-value (indicated by colour intensity) using hierarchical clustering. Associations are coloured by direction of effect (blue = positive, orange = negative), with associations significant at study-wise FDR < 0.05 marked with +/-for positive/negative correlations. Coloured associations without a label indicate nominally significant associations (p-value < 0.05, no multiple testing correction). **b**, Comparison of correlations between signatures predictive for diseases (lower triangle) and comorbidities of these diseases in the cohort (upper triangle).

To further validate the commonality of the microbiome signatures for diseases, we constructed L1/L2 regularized regression prediction models for the 36 most common diseases with > 100 cases each in the cohort using a randomly selected 90% of our data as a training set and 10% as a test set.

The microbiome-driven predictive signature of health (*lack of any reported disease*) showed a prediction area under the curve (AUC) of 0.62 on the training set and of 0.57 on the test set, which was comparable with the AUC for prediction of general anthropomorphic parameters (age, sex and BMI; AUC of 0.58 on both training/test sets). Combining microbiome and anthropometric data resulted in AUCs of 0.64 and 0.61 (training/test set). L1/L2 regression selected 73 microbial features for the “health/disease” model fit, of which the majority were microbial taxa rather than pathways (Supplementary table 6a). These features were found to be largely consistent with the healthy microbiome signature we had identified in the association analysis (Fig. 4), with 22/31 species selected by the model also associated with > 5 diseases in our analyses. The microbiome signatures for 29 out of 36 tested diseases showed AUCs > 0.55 on the test set, with the highest predictive power observed for T2D (AUC = 0.81), gallstones (AUC = 0.66) and kidney stones (AUC = 0.64). No predictive power (AUC ≤ 0.5) was observed for severe acne, undefined anaemia, non-hip fractures and reports of high cholesterol (Supplementary table 6b). Additionally, microbiome predictions of diseases showed high correlation despite the low disease comorbidities in our cohort (Fig. 4b). Finally, we calculated the recently developed Gut Microbiome Health Index (GMHI)^51^ for our data and identified a significant difference in GMHI between healthy and unhealthy individuals (p-value = 1.1e-11, balanced accuracy 0.61, Supplementary fig. 6). Our microbiome signature of health showed high overlap with signatures reported in the GMHI study^51^, with 43/50 GMHI signals replicating across studies at genus or species level (Supplementary table 1). In addition to the observed consistency in families *Clostridiaceae, Ruminococcaceae* and *Lachnospiraceae* and genera *Bacteroides, Bilophila, Coprococcus, Faecalibacterium, Ruminococcus* and *Sutterella*, we also discovered 55 novel microbiome–health associations not identified in the GMHI study, including for species from *Butyrivibrio* and *Akkermansia* and *Prevotella* genera (Supplementary tables 3b,10).

### The disease-associated microbiome is a combination of disease and medication signatures

We observed a large overlap between disease-associated microbiome patterns and microbiome associations with medication use (Supplementary fig. 7, Supplementary tables 3a-g), with the largest effects observed for proton pump inhibitors (PPIs), antibiotics, biguanide antidiabetics, osmotic laxatives and intestinal anti-inflammatory agents (84, 56, 47 and 32 associations with microbial taxa, respectively, at FDR < 0.05). To disentangle medication use from the diseases for which these drugs are prescribed, we constructed microbiome-drug-disease multivariate models for diseases where drug use is strongly correlated and/or specific for the disease: antidiabetics in T2D, selective serotonin reuptake inhibitors in depression and PPIs in functional GI disorders/IBS. We observed that both drug use and presence of the disease are independently and significantly associated with microbiome features in these models (Supplementary table 7), with consistent effect directions and strength. This observation may indicate that the unhealthy gut microbiome signature reflects both the disease and the medication used to treat it.

### Childhood environment is associated to healthy microbiome in adult life

Since it is known that the first 2–3 years of life are crucial for microbiome development, we examined the influence of early-life (age < 4 years) factors on the adult microbiome. We identified 106 associations with taxa, 30 with pathways, 22 with antibiotic resistance genes and 2 with virulence factors (FDR < 0.05), with only minimal effects observed for birth mode, breastfeeding and preterm birth (Fig. 3b, Supplementary fig. 8). Childhood living environment (defined on a scale from 1 = rural to 5 = highly urban) was significantly associated with the adult microbiome (54, 8 and 7 associations with taxa, pathways and CARDs, respectively, at FDR < 0.05). A rural childhood environment was associated with an increase in various species of bacteria, including *P. copri, F. prausnitzii, Rothia mucilaginosa* and various species of *Bifidobacterium* and *Mitsuokella* genera, abundances of which were also positively associated with increased general health. In contrast, gut abundances of species from genera *Bacteroides, Alistipes* and *Biophila* were reduced in participants with urban childhood environments.

Furthermore, parental smoking was associated with the microbiome composition of their children (15, 9 and 4 associations with taxa, pathways and CARDs, respectively, at FDR < 0.05). Here we observed associations between parental smoking and decreased abundances of *F. prausnitzii* and *P. copri*, consistent with observations for current smokers. Finally, pet ownership during childhood was associated with the adult microbiome (7 associations at FDR < 0.05), with decreases in *Alistipes finegoldii, Lactobacillus delbrueckii* and species from *Dialister* and *Bilophila* genera observed in participants who had pets as children.

We observed minimal effects for birth mode, breastfeeding and preterm birth (Fig. 3b, Supplementary fig. 8). These included an association of C-section with increases in *P. copri* in adults and of higher birth weight with increases in *Bacteroides eggerthii* and *Butyrivibrio crossotus*.

### Urbanicity, smoking, pollutants and pets are linked to health-associated bacteria

Next, we studied environmental factors at time of sampling. We identified significant shared positive association patterns between healthy microbiome signatures, pet ownership, rural living environment and greenspace surface area in the living environment (Fig. 3b), including increases in *P. copri, Bacteroides plebeius, Desulfovibrio piger* and species of genus *Mitsuokella* and decreases in *Bacteroides fragilis* and *Bilophila wadsworthia*. These associations were in contrast to the associations seen for increased measurements of NO_2_ and small particulate matter pollutants, which showed microbiome signatures in the opposite direction and negative association with health (Supplementary fig. 9).

Smoking phenotypes, including current active and passive smoking and a history of past smoking, were among the strongest phenotypes associated with microbiome composition in our study. Active smoking was associated with 41 species and 84 pathways (FDR < 0.05, Fig. 3b), with 60% of these factors also associated with previous smoking, suggesting a long-lasting signature of smoking. Intriguingly, 15 of these were further associated with passive smoking, highlighting the need to consider passive smoking in disease risk models. The directionality of associations was consistent across the three smoking phenotypes and shared direction with microbiome signatures of diseases, including decreased abundance of the butyrate-producing bacteria *Roseburia hominis, F. prausnitzii, Coprococcus catus* and *Subdoligranulum spp*; decreased abundance of the facultative oral bacteria *Veillonella spp* and *Haemophilus parainfluenzae*; and increased abundance of several species within order Clostridiales, including *Flavonifractor plautii, Pseudoflavonifractor capillosus* and *Oscillibacter spp* (Supplementary fig. 9).

### High dietary quality is linked to health-associated microbes

We identified 378 associations between 20 dietary factors, which were found to be stable over a 5-year period between food frequency questionnaire (FFQ) collection and faecal sampling (Supplementary table 8, Supplementary Discussion), and 82 species (Supplementary table 3f). The Lifelines diet score (LLDS), a diet quality score based on international nutrition literature^14^, showed the highest number of associations (79 associations with taxa, 44 with pathways, 20 with antibiotic resistance genes and 8 with virulence factors at FDR < 0.05), followed by total alcohol intake, glycemic load, protein diet score (reflecting quantity and source of protein) and total carbohydrate intake. The LLDS and protein intake scores, including animal protein, showed association patterns that overlapped with signals between microbiome features and increased general health, such as decreases in *Clostridia* species, increases in *Butyrivibrio* and *Roseburia* genera and pathways involved in ubiquinol and menaquinol synthesis. In contrast, total dietary carbohydrate intake and glycemic load showed the opposite associations (Fig. 3b, Supplementary fig. 10).

### Socio-economic factors are associated with microbiome composition

Several measurements related to the socio-economic status of the participants were available for our cohort. This included information about the number of working hours, income per month, religion and neighbourhood characteristics. In total, we observed 220 significant associations (FDR < 0.05). Of these, 72 were between bacterial abundances and monthly salary, with higher income associated with a healthy microbiome signature, such as an increase in *F. prausnitzii, Akkermansia municiphila* and *Bifidobacterium adolescentis*. In contrast, the “unhealthy microbiota” *Clostridium bolteae, Ruminococcus gnavus* and *Streptococcus parasanguinis* were more abundant in participants with lower incomes. Similar patterns were observed when analysing the differences between people living in high-income neighbourhoods versus low-income neighbourhoods. In our cohort, a high income showed low but significant correlations with neighbourhood greenspace area, rural living environment and higher LLDS (Spearman correlations 0.22, 0.17 and 0.07, respectively; correlation test FDRs < 1.0e-6), all of which share microbiome patterns observed in high income participants, implying that microbiome–income association is likely a combination of multiple factors, including a healthier diet and lifestyle and a less urban living environment.

## Discussion

Microbiome studies have associated the gut microbiome to diverse chronic and acute diseases^1–3^, to the functioning of the immune system^52^ and to drug response^53^, highlighting that the gut microbiome is an essential factor in maintenance of human health. While multiple microbiome-targeting therapies are currently being investigated in clinical trials^54^, the therapeutic potential of microbiome-targeting interventions is still confounded by a lack of consensus in the definition of a healthy or unhealthy microbiome and by a limited understanding of how heritability, exposome, lifestyle and diet shape microbiome signatures of health and disease. To bridge this gap, we analysed the microbiomes of 8,208 individuals from an extensively phenotyped three-generation cohort, allowing us to pinpoint core and keystone microbiome features and assess the effect of technical variables, anthropometrics, heritability, cohousing, diseases, diet and environmental exposures on the microbiome.

In addition to confirming known microbiome associations with age, sex and BMI^43,55^, our results highlight the importance of considering often-omitted confounders related to the stool samples (stool consistency), sampling season and sample processing (such as DNA concentration or sequencing batch). This is especially important when studying diseases where age, sex, BMI and faecal consistency are often associated with the disease.

Our observation that the microbiome is primarily associated to cohabitation and environment rather than relatedness corroborates previous studies that identified limited overall microbiome heritability^56^ and divergence in microbiomes of twins who stopped cohousing^57^. However, we do identify that the core component of the gut microbiome is significantly heritable, whereas it is the disease-associated microbes that are largely influenced by environment. These results suggest that the bacteria with low heritability that are enriched in diseases, e.g. species from genera *Clostridia, Flavonifracter* and *Veillonella*, might be more susceptible to microbiome-altering therapies than more heritable bacteria such as those from genera *Sutterella, Collinsella* and *Bacteroides*.

By comparing associations between microbiome, health and diverse diseases, we identified a common signal for dysbiosis in the gut (Fig. 4) that was largely consistent with a previous study.^51^ The existence of shared dysbiosis has considerable implications for microbiome research and development of microbiota-targeting diagnostics and therapies. It implies that the gut microbiome is a biomarker of general health, which is supported by our prediction models and by previous studies^51,58^. However, this also complicates microbiome-based diagnosis of individual diseases because single-disease models might be confounded by signals shared across unrelated diseases, and testing such models for specificity in mixed-disease cohorts will be an important step before clinical implementation. The shared microbiome signature is also exciting because it suggests that microbiome-targeting interventions could improve overall human health. This is supported by our observations that lifestyle factors generally considered healthy, such as adherence to current dietary recommendations and no smoking, associate with similar microbiome patterns to those associated with general health. While microbiome–drug interactions are well described *in vitro*^59^, and characterized *in vivo* for antibiotics, PPIs^60^ and antidiabetics^61^, our results suggest that the general microbiome dysbiosis is a combination of drug- and disease-effects, implying that many currently understudied drugs, such as SSRIs, have a negative effect on the gut microbiome. This also highlights the importance of controlling for medication use.

By linking (un)healthy microbiome patterns to childhood and current exposome, diet and socioeconomics, we observed that a healthier diet^14^, childhood and current exposure to rural environment and pets, exposure to greenspace and higher income share signals with healthy microbiome patterns. These observations support the microbiome diversity hypothesis (also known as the hygiene hypothesis) – a postulate that reduction in exposure to microbiota contributes to an increase in the frequency of autoimmune and allergic diseases^62,63^. Notably, while the classic hygiene hypothesis focuses on pathogens and the impact of early-life exposures, our results suggest that exposures in adulthood also contribute to (un)healthy microbiome patterns, implying that the environment shapes the microbiome throughout human life and, as such, microbiome-targeted therapies could be effective throughout an individual’s life. Furthermore, while we identified negative correlations of diet scores, pets and rural environment with opportunistic pathogens, such as *Clostridia* species, we also observed positive correlation with commensals such as butyrate producers from genera *Bacteroides, Alistipes* and *Faecalibacterium*, implying that exposure not only to pathogens but also to commensals from the environment plays an important role in establishing a healthy gut ecosystem.

We also observed that smoking, a high-carbohydrate diet and exposure to NO_2_ and small particulate matter (PM_2.5_) are positively correlated with disease-linked bacterial species from genera *Clostridia* and *Ruminococcus*. While air pollutants were previously associated to GI diseases in humans^64^, and have been shown to affect the gut microbiota of mice, the effects of air pollutants on the human gut microbiota are still unclear^65^. Our results suggest that air pollutants negatively impact the human gut microbiota and might increase risk for development of GI diseases by contributing to general microbiome dysbiosis.

In addition to identifying links between current exposome and microbiome, we also found that childhood exposures to smoking, pets and rural environment are associated with the subsequent adult microbiome. While the effect sizes for these associations were lower than those of equivalent current exposures measured alongside faecal sampling, the effect directions and patterns were consistent, suggesting that environmental exposures can have a long-lasting effect on the gut microbiome. This is further supported by our observation that smokers who stopped smoking still showed microbiome associations similar to current smokers, albeit with lower effect sizes. Notably, we observed limited associations between microbiome and exposures during and shortly after birth, including breastfeeding, birth mode and preterm birth. These exposures are known to shape infant microbiota^66,67^, but our findings indicate that their effects on the adult microbiome are indiscernible, possibly because the selective pressures induced by these exposures end early in human life and these early effects are superseded over time by other selective pressures such as diet, exposure to microbiota of family members and environment.

Notably, while we measured 241 phenotypes from a broad set of categories, we could only explain ∼15% of the variation in microbiome composition and function between individuals. While this level of explained variance is consistent with those found in previous large-scale studies of European and American populations^6,56,66^, it implies that the gut microbiome is highly individual and that our current understanding of the factors that shape it is still limited. A potential explanation for the existence of this “missing variance” is that the microbiome composition and function are a result of an individual’s history of lifestyle and exposures, and cross-sectional measurement is thus insufficient to fully explain it. This is supported by our observation that early-life exposures are associated with microbiome in adult age and that cohousing participants have significantly closer microbiomes than non-cohabiting individuals, regardless of relatedness. Future quantification of this missing variance, potentially by long-time-frame longitudinal studies, will play a critical part in future development of microbiome-targeting diagnostics and therapies.

## Conclusion

We generated and analysed the largest, multi-generational gut microbiome cohort to date that has been collected and profiled in a highly standardized matter, and linked it to extensive phenotype data. We defined and described a gut dysbiosis shared across diverse diseases and identified novel links between this dysbiosis and heritability, childhood and current exposome, lifestyle and socioeconomics. This study demonstrates the power of large-scale, well-phenotyped cohorts for dissecting the links between gut microbiome, health, genetics and environment and provides a rich resource for future studies for microbiome-directed interventions.

## Supporting information

Supplementary Discussion

## Acknowledgements

We would like to acknowledge and thank the late Marten Hofker who had the great wisdom and vision to initiate the Lifelines DAG3/Dutch Microbiome Project.

The authors wish to acknowledge the services of the Lifelines Cohort Study, the contributing research centres delivering data to Lifelines and all the study participants. The Lifelines Biobank initiative has been made possible by subsidy from the Dutch Ministry of Health, Welfare and Sport; the Dutch Ministry of Economic Affairs; the University Medical Center Groningen (UMCG the Netherlands); the University of Groningen and the Northern Provinces of the Netherlands. We would like to thank the Center for Information Technology of the University of Groningen (RUG) for their support and for providing access to the Peregrine high performance computing cluster and the Genomic Coordination Center (UMCG and RUG) for their support and for providing access to Calculon and Boxy high performance computing clusters, and the MOLGENIS team for data management and analysis support. Metagenomics library preparation and sequencing was done at Novogene. We also thank K. Mc Intyre for English and content editing.

## Author contribution

RG designed and implemented the metagenomic data analysis pipelines, analysed metagenomic data, performed heritability analysis and drafted the manuscript. AK designed the prediction models and implemented statistical methods for association analyses and assisted in drafting of the manuscript. AVV, LC, VC, SH, MAYK, SAS, JB, LAB, VL, TS, MH and SS assisted in other statistical analyses, interpretation of data and drafting of the manuscript. MS provided data stewardship and analysis infrastructure. BHJ, JAMD and JGA collected data, assisted in study planning and critically reviewed the manuscript. SS supervised and coordinated heritability analysis. RCHV provided the air pollution data and supervised the air pollution analysis. HJMH, SZ, RKW, JF and CW conceived, coordinated and supported the study. All authors critically revised and approved the manuscript.

## Funding

Sequencing of the cohort was funded by a grant from the CardioVasculair Onderzoek Nederland grant (CVON 2012-03) to MH, JF and AZ. RG, HH and RKW are supported by the collaborative TIMID project (LSHM18057-SGF) financed by the PPP allowance made available by Top Sector Life Sciences & Health to Samenwerkende Gezondheidsfondsen (SGF) to stimulate public-private partnerships and co-financing by health foundations that are part of the SGF. RKW is supported by the Seerave Foundation and the Dutch Digestive Foundation (16-14). AZ is supported by European Research Council (ERC) Starting Grant 715772, Netherlands Organization for Scientific Research (NWO) VIDI grant 016.178.056, CVON grant 2018-27, and NWO Gravitation grant ExposomeNL 024.004.017. JF is supported by NWO Gravitation grant Netherlands Organ-on-Chip Initiative (024.003.001) and CVON grant 2018-27. CW is further supported by an ERC advanced grant (ERC-671274) and an NWO Spinoza award (NWO SPI 92-266). LC is supported by a joint fellowship from the University Medical Center Groningen and China Scholarship Council (CSC201708320268) and a Foundation De Cock-Hadders grant (20:20-13). MS is supported by Netherlands Organization for Scientific Research (NWO) VIDI grant 016 and EUCAN-connect, a project funded by European Commission H2020 grant 824989.

## Competing interests declaration

Authors declare no conflict of interest.

## Scripts and availability of data

Scripts used for data analysis can be found at: https://github.com/GRONINGEN-MICROBIOME-CENTRE/Groningen-Microbiome/tree/master/Projects/DMP

## Notes

### Competing Interest Statement

The authors have declared no competing interest.

## References

1. Lynch, S. V. & Pedersen, O. The Human Intestinal Microbiome in Health and Disease. N. Engl. J. Med. 375, 2369–2379 (2016).

2. Liang, D., Leung, R. K.-K., Guan, W. & Au, W. W. Involvement of gut microbiome in human health and disease: brief overview, knowledge gaps and research opportunities. Gut Pathog. 10, 3 (2018).

3. Duvallet, C., Gibbons, S. M., Gurry, T., Irizarry, R. A. & Alm, E. J. Meta-analysis of gut microbiome studies identifies disease-specific and shared responses. Nat. Commun. 8, 1784 (2017).

4. Zmora, N., Soffer, E. & Elinav, E. Transforming medicine with the microbiome. Sci. Transl. Med. 11, (2019).

5. Zmora, N., Zeevi, D., Korem, T., Segal, E. & Elinav, E. Taking it Personally: Personalized Utilization of the Human Microbiome in Health and Disease. Cell Host Microbe 19, 12–20 (2016).

6. Zhernakova, A. et al. Population-based metagenomics analysis reveals markers for gut microbiome composition and diversity. Science 352, 565–569 (2016).

7. Gaulke, C. A. & Sharpton, T. J. The influence of ethnicity and geography on human gut microbiome composition. Nat. Med. 24, 1495–1496 (2018).

8. Vatanen, T. et al. Genomic variation and strain-specific functional adaptation in the human gut microbiome during early life. Nat. Microbiol. 4, 470–479 (2019).

9. Scholtens, S. et al. Cohort Profile: LifeLines, a three-generation cohort study and biobank. Int. J. Epidemiol. 44, 1172–1180 (2015).

10. Tigchelaar, E. F. et al. Cohort profile: LifeLines DEEP, a prospective, general population cohort study in the northern Netherlands: study design and baseline characteristics. BMJ Open 5, (2015).

11. Siebelink, E., Geelen, A. & de Vries, J. H. M. Self-reported energy intake by FFQ compared with actual energy intake to maintain body weight in 516 adults. Br. J. Nutr. 106, 274–281 (2011).

12. Brouwer-Brolsma, E. M. et al. A National Dietary Assessment Reference Database (NDARD) for the Dutch Population: Rationale behind the Design. Nutrients 9, (2017).

13. Willett, W. C. et al. Reproducibility and validity of a semiquantitative food frequency questionnaire. Am. J. Epidemiol. 122, 51–65 (1985).

14. Vinke, P. C. et al. Development of the food-based Lifelines Diet Score (LLDS) and its application in 129,369 Lifelines participants. Eur. J. Clin. Nutr. 72, 1111–1119 (2018).

15. G, M. et al. A Protein Diet Score, Including Plant and Animal Protein, Investigating the Association with HbA1c and eGFR-The PREVIEW Project. Nutrients vol. 9 https://pubmed.ncbi.nlm.nih.gov/28714926/ (2017).

16. Leeming, E. R., Johnson, A. J., Spector, T. D. & Le Roy, C. I. Effect of Diet on the Gut Microbiota: Rethinking Intervention Duration. Nutrients 11, (2019).

17. Eeftens, M. et al. Development of Land Use Regression Models for PM2.5, PM2.5 Absorbance, PM10 and PMcoarse in 20 European Study Areas; Results of the ESCAPE Project. https://pubs.acs.org/doi/full/10.1021/es301948k (2012) xdoi:10.1021/es301948k.

18. Development of NO2 and NOx land use regression models for estimating air pollution exposure in 36 study areas in Europe – The ESCAPE project. Atmos. Environ. 72, 10–23 (2013).

19. Eeftens, M. et al. Stability of measured and modelled spatial contrasts in NO2 over time. Occup. Environ. Med. 68, 765–770 (2011).

20. StatLine. https://opendata.cbs.nl/#/CBS/en/.

21. Ford, A. C. et al. Validation of the Rome III Criteria for the Diagnosis of Irritable Bowel Syndrome in Secondary Care. Gastroenterology 145, 1262-1270.e1 (2013).

22. Angulo, P. et al. The NAFLD fibrosis score: A noninvasive system that identifies liver fibrosis in patients with NAFLD. Hepatology 45, 846–854 (2007).

23. Bedogni, G. et al. The Fatty Liver Index: a simple and accurate predictor of hepatic steatosis in the general population. BMC Gastroenterol. 6, 33 (2006).

24. Imhann, F. et al. The 1000IBD project: multi-omics data of 1000 inflammatory bowel disease patients; data release 1. BMC Gastroenterol. 19, 5 (2019).

25. McIver, L. J. et al. bioBakery: a meta’omic analysis environment. Bioinformatics 34, 1235–1237 (2018).

26. Langmead, B. & Salzberg, S. L. Fast gapped-read alignment with Bowtie 2. Nat. Methods 9, 357–359 (2012).

27. Truong, D. T. et al. MetaPhlAn2 for enhanced metagenomic taxonomic profiling. Nat. Methods 12, 902–903 (2015).

28. Franzosa, E. A. et al. Species-level functional profiling of metagenomes and metatranscriptomes. Nat. Methods 15, 962–968 (2018).

29. Swertz, M.A. et al. The MOLGENIS toolkit: rapid prototyping of biosoftware at the push of a button. BMC Bioinformatics 11 S12 (2010).

30. Hsieh, T. C., Ma, K. H. & Chao, A. iNEXT: an R package for rarefaction and extrapolation of species diversity (Hill numbers). Methods Ecol. Evol. 7, 1451–1456 (2016).

31. Kaminski, J. et al. High-Specificity Targeted Functional Profiling in Microbial Communities with ShortBRED. PLoS Comput. Biol. 11, (2015).

32. Chen, L. et al. VFDB: a reference database for bacterial virulence factors. Nucleic Acids Res. 33, D325–328 (2005).

33. Jia, B. et al. CARD 2017: expansion and model-centric curation of the comprehensive antibiotic resistance database. Nucleic Acids Res. 45, D566–D573 (2017).

34. Pilia, G. et al. Heritability of Cardiovascular and Personality Traits in 6,148 Sardinians. PLoS Genet. 2, (2006).

35. Chen, W.-M. & Abecasis, G. R. Estimating the power of variance component linkage analysis in large pedigrees. Genet. Epidemiol. 30, 471–484 (2006).

36. Pincus, R. Aitchison, J.: The Statistical Analysis of Compositional Data. Chapman and Hall, London & New York 1986, XII, 416 pp. Biom. J. 30, 794–794 (1988).

37. Friedman, J. & Alm, E. J. Inferring Correlation Networks from Genomic Survey Data. PLOS Comput. Biol. 8, e1002687 (2012).

38. Chen, L. et al. Gut microbial co-abundance networks show specificity in inflammatory bowel disease and obesity. Nat. Commun. 11, 1–12 (2020).

39. M, A. et al. Enterotypes of the human gut microbiome. Nature vol. 473 https://pubmed.ncbi.nlm.nih.gov/21508958/ (2011).

40. Berry, D. & Widder, S. Deciphering microbial interactions and detecting keystone species with co-occurrence networks. Front. Microbiol. 5, 219 (2014).

41. J, Q. et al. A human gut microbial gene catalogue established by metagenomic sequencing. Nature vol. 464 https://pubmed.ncbi.nlm.nih.gov/20203603/ (2010).

42. Turnbaugh, P. J. et al. The Human Microbiome Project. Nature 449, 804–810 (2007).

43. T, Y. et al. Human gut microbiome viewed across age and geography. Nature vol. 486 https://pubmed.ncbi.nlm.nih.gov/22699611/ (2012).

44. Jk, G. et al. Human genetics shape the gut microbiome. Cell vol. 159 https://pubmed.ncbi.nlm.nih.gov/25417156/ (2014).

45. Faecalibacterium prausnitzii and human intestinal health. Curr. Opin. Microbiol. 16, 255–261 (2013).

46. Eppinga, H. et al. Similar Depletion of Protective Faecalibacterium prausnitzii in Psoriasis and Inflammatory Bowel Disease, but not in Hidradenitis Suppurativa. J. Crohns Colitis 10, 1067–1075 (2016).

47. R, H. et al. Metagenome-wide association study of the alterations in the intestinal microbiome composition of ankylosing spondylitis patients and the effect of traditional and herbal treatment. Journal of medical microbiology vol. 69 https://pubmed.ncbi.nlm.nih.gov/31778109/ (2020).

48. Kandeel, W. A. et al. Impact of Clostridium Bacteria in Children with Autism Spectrum Disorder and Their Anthropometric Measurements. J. Mol. Neurosci. 70, 897–907 (2020).

49. Tap, J. et al. Identification of an Intestinal Microbiota Signature Associated With Severity of Irritable Bowel Syndrome. Gastroenterology 152, 111-123.e8 (2017).

50. Vieira-Silva, S. et al. Statin therapy is associated with lower prevalence of gut microbiota dysbiosis. Nature 581, 310–315 (2020).

51. Gupta, V. K. et al. A predictive index for health status using species-level gut microbiome profiling. Nat. Commun. 11, 1–16 (2020).

52. Thaiss, C. A., Zmora, N., Levy, M. & Elinav, E. The microbiome and innate immunity. Nature 535, 65–74 (2016).

53. Precision Medicine Goes Microscopic: Engineering the Microbiome to Improve Drug Outcomes. Cell Host Microbe 26, 22–34 (2019).

54. Garber, K. First microbiome-based drug clears phase III, in clinical trial turnaround. Nature Reviews Drug Discovery vol. 19 655–656 https://www.nature.com/articles/d41573-020-00163-4 (2020).

55. Castaner, O. et al. The Gut Microbiome Profile in Obesity: A Systematic Review. International Journal of Endocrinology vol. 2018 e4095789 https://www.hindawi.com/journals/ije/2018/4095789/ (2018).

56. Rothschild, D. et al. Environment dominates over host genetics in shaping human gut microbiota. Nature 555, 210–215 (2018).

57. Shotgun Metagenomics of 250 Adult Twins Reveals Genetic and Environmental Impacts on the Gut Microbiome. Cell Syst. 3, 572-584.e3 (2016).

58. Oh, M. & Zhang, L. DeepMicro: deep representation learning for disease prediction based on microbiome data. Sci. Rep. 10, 1–9 (2020).

59. M, Z., M, Z.-K., R, W. & Al, G. Mapping human microbiome drug metabolism by gut bacteria and their genes. Nature vol. 570 https://pubmed.ncbi.nlm.nih.gov/31158845/ (2019).

60. Imhann, F. et al. Proton pump inhibitors affect the gut microbiome. Gut 65, 740–748 (2016).

61. Forslund, K. et al. Disentangling type 2 diabetes and metformin treatment signatures in the human gut microbiota. Nature 528, 262–266 (2015).

62. Bach, J.-F. The hygiene hypothesis in autoimmunity: the role of pathogens and commensals. Nat. Rev. Immunol. 18, 105–120 (2018).

63. Scudellari, M. News Feature: Cleaning up the hygiene hypothesis. Proc. Natl. Acad. Sci. 114, 1433–1436 (2017).

64. Salim, S. Y., Kaplan, G. G. & Madsen, K. L. Air pollution effects on the gut microbiota. Gut Microbes (2013) doi:10.4161/gmic.27251.

65. Impact of air quality on the gastrointestinal microbiome: A review. Environ. Res. 186, 109485 (2020).

66. Manor, O. et al. Health and disease markers correlate with gut microbiome composition across thousands of people. Nat. Commun. 11, 1–12 (2020).

