## Supplementary Discussion for "The Dutch Microbiome Project defines factors that shape the healthy gut microbiome"

### Supplementary Discussion: Association of gut microbiome to diseases and medication use

When studying gut microbiome signatures in 82 measurements of health and diseases, we identified that a total of 1,206 significant associations with taxa, 1,182 with pathways, 390 with antibiotic resistance gene families and 76 with bacterial virulence factors (FDR < 0.05, Supplementary tables 3B-3F). Examined metadata included self-reported health (RAND questionnaire reports of self-reported mental and physical health, RAND questionnaire health change in previous year, general health questionnaire report of no serious diseases/disorders and general health questionnaire history of appendectomy) as well as 78 diseases collected via general health questionnaire (Supplementary tables 3A-3H).

***Microbiome-disease associations are dominated by gastrointestinal disorders***

Among disease groups, we observed the strongest signal for GI diseases (386 associations with taxa and 433 with pathways at FDR<0.05) involving disorders such as inflammatory bowel disease, irritable bowel syndrome and non-alcoholic fatty liver disease. Participants experiencing GI discomfort (self reported IBS), displayed the largest amount of associations with 48 different taxa, and a decrease in α-diversity. Shared gut microbiome signatures in GI-diseases consisted mainly of decreases in *Faecalibacterium prausnitzii*, *Subdoligranulum unclassified*, *Eubacterium rectale, Bifidobacterium adolescentis* and *Alistipes senegalensis*, and increases in *Ruminococcus gnavus*, *Clostridium asparagiforme*, *Clostridium bolteae*, *Flavonifractor plautii*, *Eggerthella unclassified* and *Clostridium citroniae*. In addition, decreases were observed in genes encoding pathways involved in the degradation of glutaryl-CoA, deoxy-l-threo-hex-4-enopyramironate, fructuronate and hexuronate, the salvage of adenosylcobalamin from cobalamide and the biosynthesis of putrescine. Moreover, increases were observed in microbial genes encoding pathways involved in the biosynthesis of kdo-2-lipid-A, ubiquinol-8, enterobacterial common antigen and L-tryptophan, the degradation of L-arginine, and in sulfo glycolysis. More specifically, increases in *Ruminococcus gnavus*, *Clostridium asparagiforme*, *Clostridium bolteae*, *Flavonifractor plautii* and *Eggerthella unclassified* were observed in participants affected by GI discomfort (diarrea, nausea, feeling unwell, stomach ache), irritable bowel syndrome and gallstones. *Clostridium bolteae* was also increased in participants affected by Crohn's disease (CD), ulcerative colitis (UC) and in bloating and diarrhea. Moreover, *Clostridium* *asparagiforme* was increased in participants with GI discomfort (bloating, burping, diarrhea), celiac disease and hepatitis. Microbial genes encoding pathways involved in the degradation of glutaryl-CoA,deoxy-l-threo-hex-4-enopyramironate and the salvage of adenosylcobalamin from cobamamide were decreased in participants affected by hepatitis, gallstones, GI-discomfort (nausea, unwell, pain, diarrea, bloating, constipation), CD, IBS and stomach ulcers.

Of the medication used for diseases included in this GI-category, usage of PPIs (ATC-code A02BC), laxatives (ACT-code A06AD) and antibiotics in the 3 months prior to sampling, were associated with the most taxa (84, 72, and 117 associations at FDR < 0.05) and pathways (196, 205, 74 associations at FDR < 0.05) in the gut microbiome. Interestingly, PPIs usage was associated with increases in *Streptococcus parasanguinis* and *Streptococcus salivarius* which were also associated with stomach ulcers and feeling bloated. Moreover, the gut microbiomes of PPI users displayed a decrease in genes encoding the biosynthesis of amino acids and the fermentation of pyruvate to acetate and lactate, which was also observed in participants having a stomach ulcer. Laxative use was associated with an increase in *Veillonella parvula*, *Ruminococcus torques* and *Streptococcus parasanguinis* and with numerous pathways, but these species and pathways were not associated with having constipation.

***Cardiologic, metabolic and haematologic diseases***

When exploring gut microbiome signatures of cardiologic, metabolic and haematologic origin, we identified a total of 484 associations between 123 unique taxa and 217 microbial pathways with 14 diseases (FDR<0.05). We observed an increase of *Streptococcus parasanguinis* in participants affected by type 2 diabetes, arrhythmias, an increased heart rate, atherosclerose, a heart attack and heart failure. Moreover, *Streptococcus salivarius* and *Veilonella unclassified* were increased in participants affected by arrhythmias, a heart attack and heart failure. In addition, increases of *C. bolteae*, *F. plautii* and *C. asparagiforme* were observed in participants affected by type 2 diabetes and hypertension. Furthermore, decreases of *Alistipes senegalensis* were observed in participants affected by type 2 diabetes, arrhythmias, an increased heart rate, heart valve problems, high cholesterol and hypertension. Participants affected by type 2 diabetes, a heart attack or high cholesterol shared a decrease in genes encoding 28 pathways which were involved, among others, in the formation and salvage of thiamin (vitamin B1), amino acid biosynthesis, homolactic fermentation and fermentation of pyruvate to acetate and lactate, and de novo biosynthesis of deoxyribonucleotides.

Of the medication used for cardiologic, metabolic and haematologic diseases, biguanides (ATC-code A10BA), sulfonylurea derivatives (ATC-code A10BB), beta-blockers (ATC-code C07AB), calcium antagonists (ATC-code C08CA) and statines (ATC-code C10AA) had the highest number of associations with 73 unique taxa and 241 unique pathways in the gut microbiome (132 and 766 associations at FDR < 0.05 respectively). Interestingly, a decrease in *Roseburia hominis*, *Alistipes senegalis*, and *Dorea longicatena* and an increase in *Clostridium bolteae* and *Clostridium hathewayi* were observed in participants who were using biguanides and sulfonylurea derivatives, which were also associated with having type 2 diabetes.

***Psychiatric and neurological diseases***

When exploring gut microbiome signatures of psychiatric and neurological diseases, we identified a total of 290 associations containing 90 unique taxa and 92 pathways, with 20 diseases (FDR < 0.05). Shared gut microbiome signatures of psychiatric and neurological diseases mainly consisted of a decrease in *Faecalibacterium prausnitzii*, *Bifidobacterium adolescentis*, *Dorea longicatena* and *Subdoligranulum unclassified*. More specifically, we found an increase in *Clostridium asparagiforme* and *Anaerotrunchus colohominis* common to burnout, eating disorders and depression, headaches and fibromyalgia. No pathways were shared amongst participants affected by different neurological or psychiatric disorders, nor were many specific diseases within this category associated with the abundances of many pathways. Strikingly, having an eating disorder was associated to the largest number of pathways, namely, in these participants, an increase in genes encoding 10 pathways were observed (amongst others involved in kdo-2-lipid-A biosynthesis and short chain fatty acid degradation), as well as a decrease in 60 pathways (amongst others involved in thiamin (B1) salvage and formation, and the biosynthesis of deoxyribonucleotide and amino acids).

#### *Microbiome signatures are shared across disease groups*

After exploring the gut microbiome in diseases specific to single tracti, we moved into gut microbiome signatures that are shared across diseases of different tracti to identify signatures that are possible indicative of not being healthy. Shared gut microbiome signatures across diseases of different etiology mainly consisted of increases in species from *Anaerotruncus*, *Ruminococcus*, *Bacteroides*, *Holdemania*, *Flavonifractor*, *Eggerthella* and *Clostridium* genera, and of decreases in *Faecalibacterium*, *Bifidobacterium*, *Butyrivibrio*, *Subdoligranulum*, *Oxalobacter*, *Eubacterium* and *Roseburia*. More specifically, increases of *Anaerotruncus*, *Flavonifractor* and *Clostridium* were observed in participants affected by type 2 diabetes, diarrhea and constipation, mental disorders and gallstones. Moreover, increases of *Bifidobacterium*, *Butyrivibrio* and *Subdoligranulum* were observed in participants affected by type 2 diabetes, diarrhea and chronic obstructive pulmonary disorder and irritable bowel syndrome. In addition, gut microbiome pathways that were shared across diseases of different tracti mainly consisted of increases in the biosynthesis of L-ornithine, ubiquinol and menaquinol, enterobacterial common antigen, Kdo-2-lipid-A, and molybdenum cofactor, and of decreases in the biosynthesis of amino acids, deoxyribonucleosides and nucleotides, in anaerobic energy metabolism and the fermentation to short chain fatty acids (mainly butanoate). More specifically, genes encoding the biosynthesis of ubiquinol-8 and enterobacterial common antigen and the degradation of L-arginine and methylglyoxal were increased in the gut microbiomes of participants affected by diarrhea, gallstones, chronic obstructive pulmonary disorders (COPD), and Crohn’s disease. Moreover, genes encoding several thiamin (B1) salvation and formation pathways were decreased in the gut microbiomes of participants affected by gallstones, COPD, asthma, hypertension, rheumatoid arthritis, hepatitis, eating disorder, heart attack, high cholesterol and GI discomfort.

#### *Self-reported health status reflects microbiome patterns shared across diseases*

Interestingly, participants that reported to have no diseases, had the second highest number of associations (72 significant associations with taxa, 21 with pathways), mainly displaying an increase in *Prevotella copri*, *Desulfovibrio piger*, *Faecalibacterium prausnitzii*, *Roseburia hominis*, *Bifidobacterium catenulatum*, *Dorea longicatena*, *Alistipes shahii* and

*Alistipes putredinis*, and a decrease in *Clostridium bolteae*, *Flavonifractor plautii*, H*oldemania filiformis*, *Ruminococcus gnavus*, and *Eggerthella unclassified*. Moreover, participants that reported not to have any diseases displayed an increase in their gut microbiome α-diversity. These participants also displayed an increase in gut microbial genes encoding pathways involved in the biosynthesis of putrescine, amino acids, the degradation of galactose glutaryl-CoA and fructuronate, and in the fermentation of pyruvate into acetate and lactate, as well as a decrease in genes encoding the biosynthesis of enterocommon bacterial antigen.

### Supplementary discussion: diet stability over 5-year time period

Since FFQs for all participants were recorded 5 years prior to the fecal sampling, whereas FFQ at the fecal sampling was available for a subset of items, we compared the intake of major food groups between two timepoints in a full Lifelines dataset which includes 128,501 individuals, as described in methods. We observed > 97% consistency in the intake of major food groups and dietary habits between these FFQs and more limited FFQs collected at time of sampling (Supplementary table 8), implying that the diet of study participants was largely stable over the 5 year time period prior to sampling.
