## Supplementary Figures for "The Dutch Microbiome Project defines factors that shape the healthy gut microbiome"

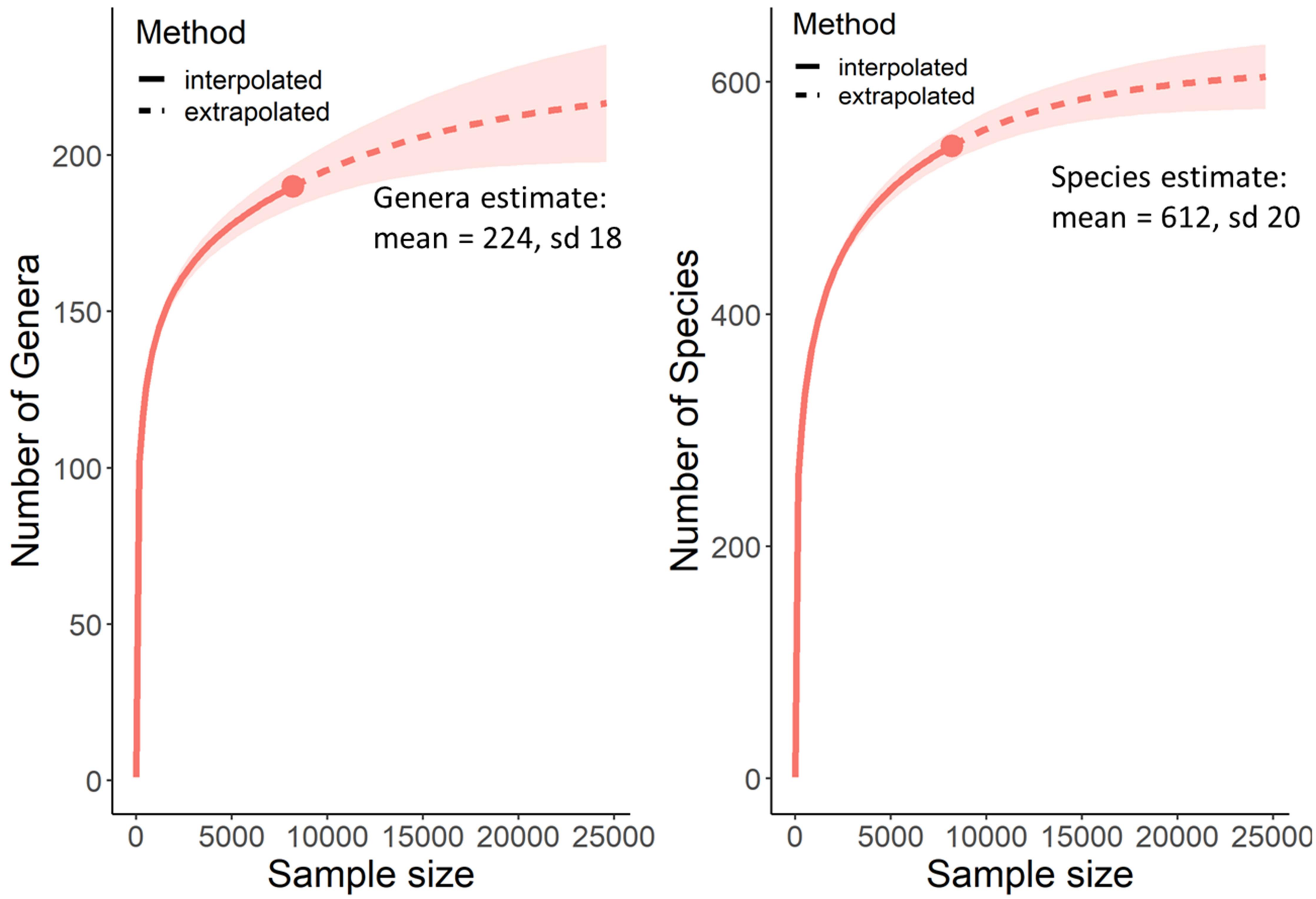


***Supplementary figure 1: Estimation of total number of species and genera in the DMP population***

*Figure shows rarefaction and extrapolation sampling curve for species and genera richness calculated using Hill numbers implemented in iNEXT package for R. Extrapolated part of rarefaction curve is shown dotted. Standard deviation of the estimate is shaded, and asymptotic richness estimate is shown on the plots.*

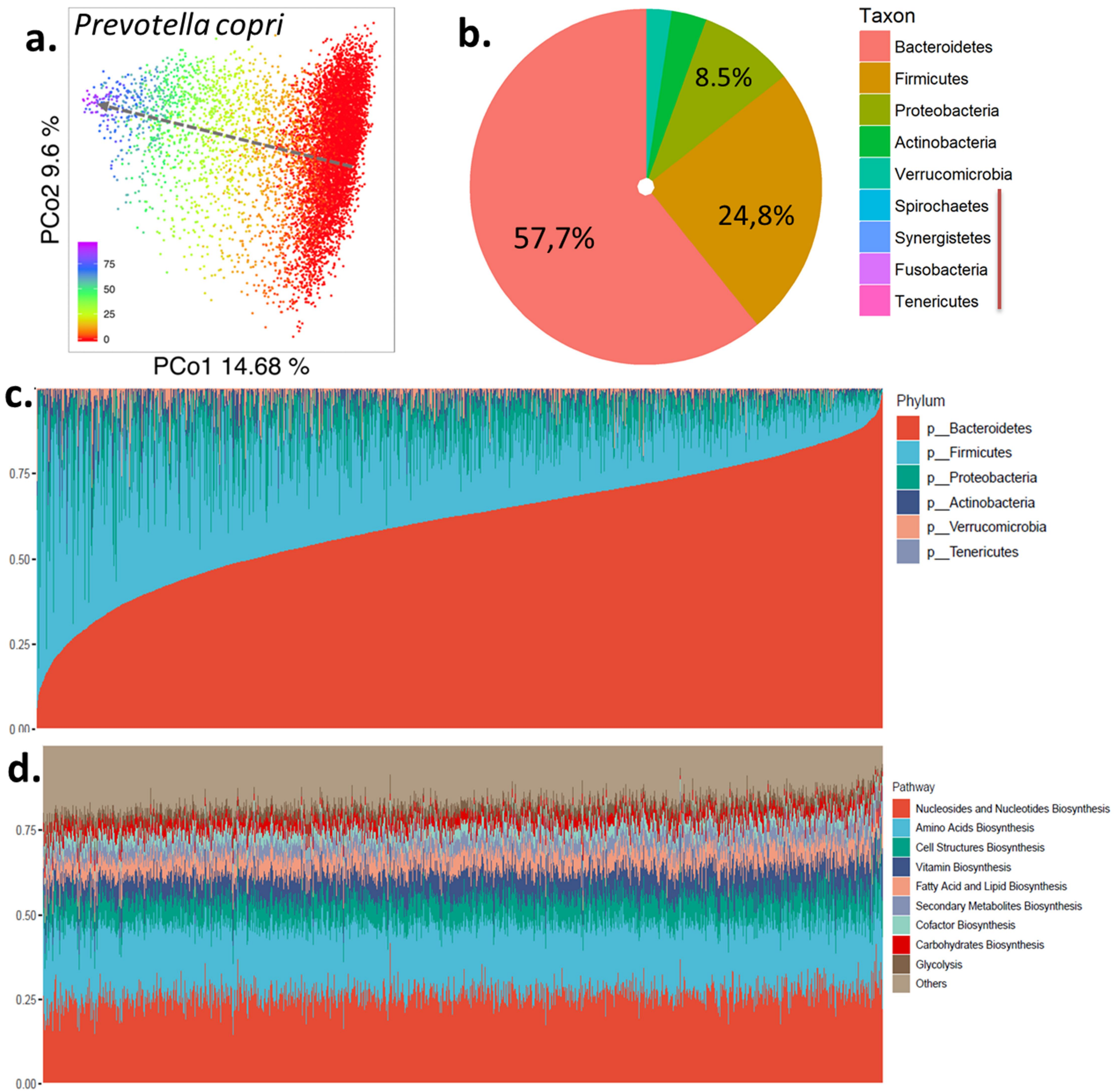


***Supplementary figure 2: Overview of DMP microbiome composition and function***

***a,** First two principal coordinates of the Bray-Curtis distance matrix calculated on microbial species of DMP cohort, colored by the relative abundance of Prevotella copri bacterium. **b,** Average relative abundances of bacterial phyla in the DMP cohort. **c,** Phylum-level composition of all samples in the cohort, sorted by abundance of phylum Bacteroidetes, with samples displayed as vertical lines. **d,** Relative abundances of top ten MetaCyc pathways of all samples, with samples displayed as vertical lines.*

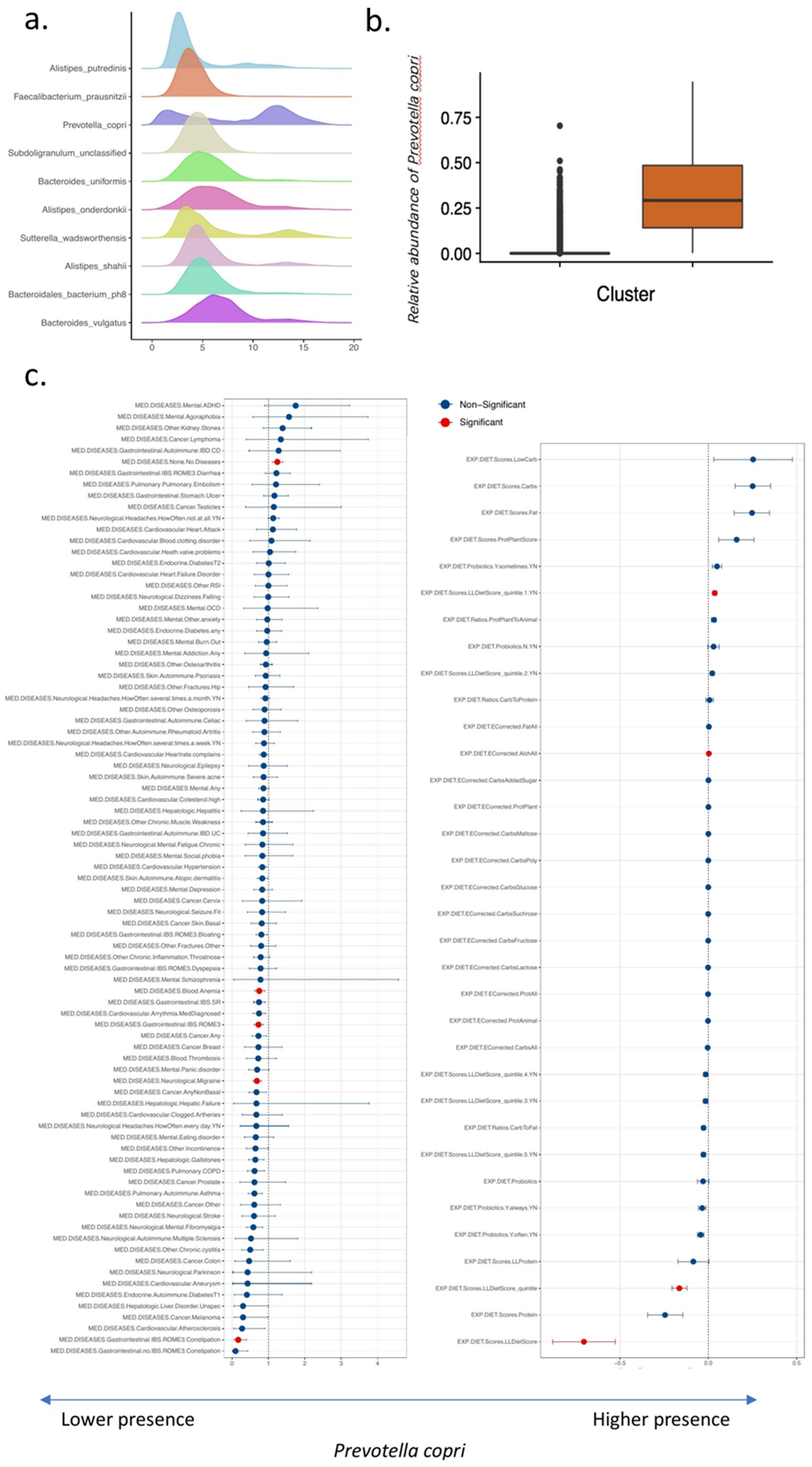


***Supplementary figure 3: Clusters determined by bi-modally distributed Prevotella copri***

***a,*** *Density plots of top ten bacterial species by relative abundance.* ***b,*** *First two principal coordinates of the Bray-Curtis distance matrix calculated on microbial species of DMP cohort, colored clusters assigned based on relative abundance of P. copri.* ***c.*** *Association of P. copri with metadata.*

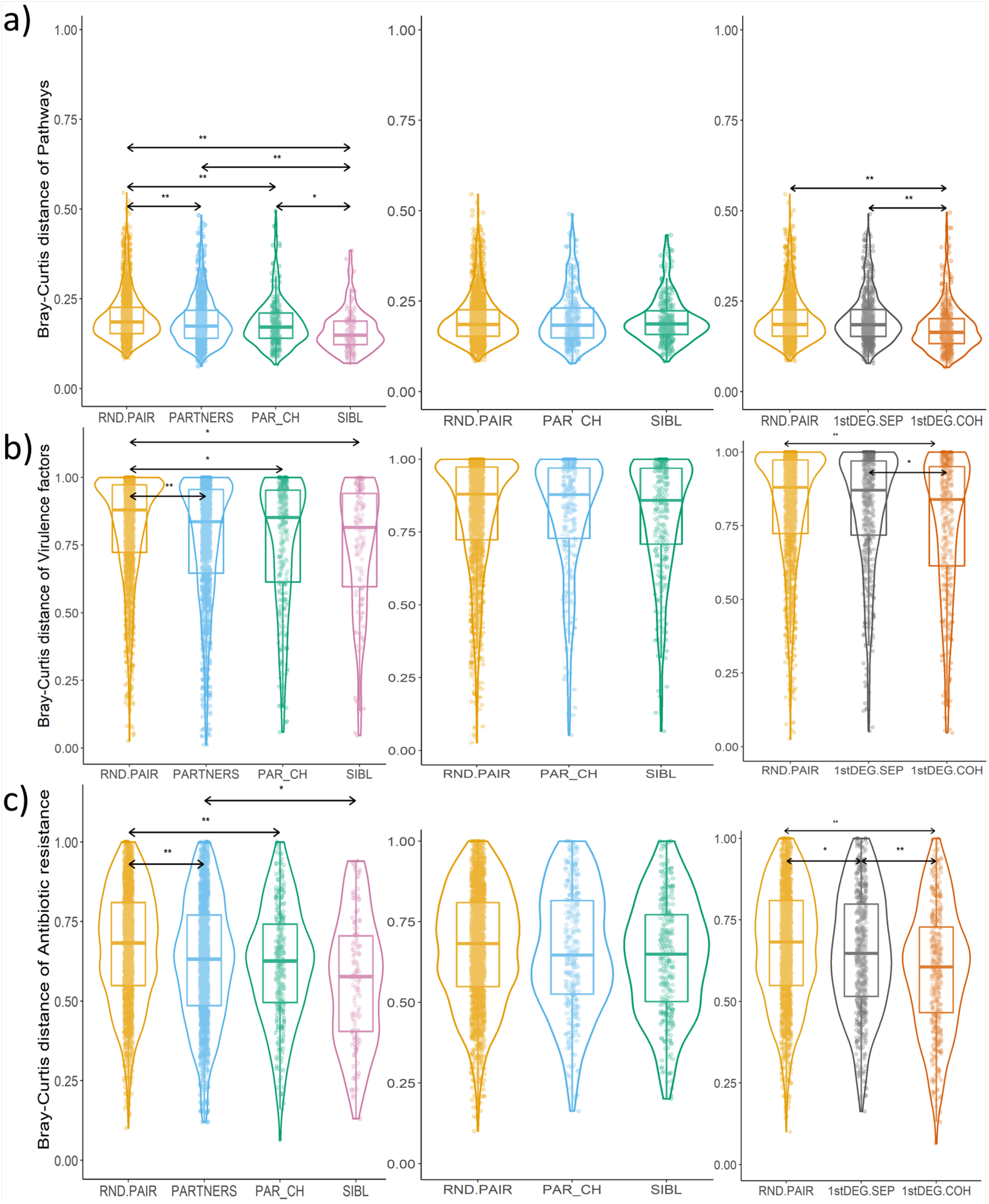


***Supplementary figure 4: Bray-curtis distances of microbiome features of cohabiting and non-cohabiting participants***

*Pairwise microbiome Bray-Curtis dissimilarity comparisons of groups of random, non-cohabiting, pairs (RND.PAIR) compared to cohabitating partners (PARTNERS), cohabiting parent-child pairs (PAR_CH) and cohabiting siblings (SIBL); non-cohabiting random pairs, parent-child pairs and sibling pairs; and random pairs compared to non-cohabiting 1$^{st}$ degree relatives (1stDEG.SEP) and cohabiting 1$^{st}$ degree relatives (1stDEG.COH).* ***a,*** *MetaCyc pathways,* ***b,*** *Virulence factor gene families, and* ***c,*** *antibiotic resistance gene families. Significantly different groups are marked with ** for FDR < 1.0e-5 or * for FDR < 0.05.*

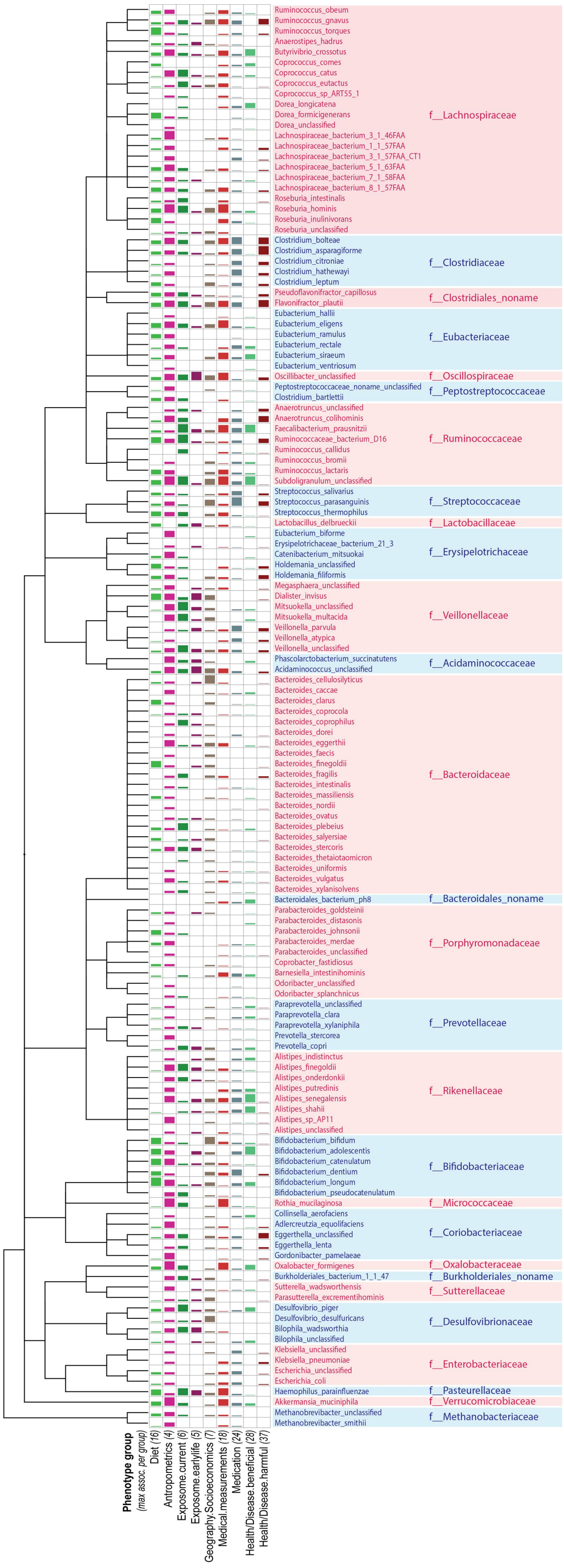


***Supplementary figure 5: Overview of microbiome-phenotype associations***

*Figure shows a number of study-wide significant associations (FDR < 0.05) per phenotype group, clustered by taxonomy, with bar sizes representing the number of associations relative to maximal number of associations for the phenotype group.*

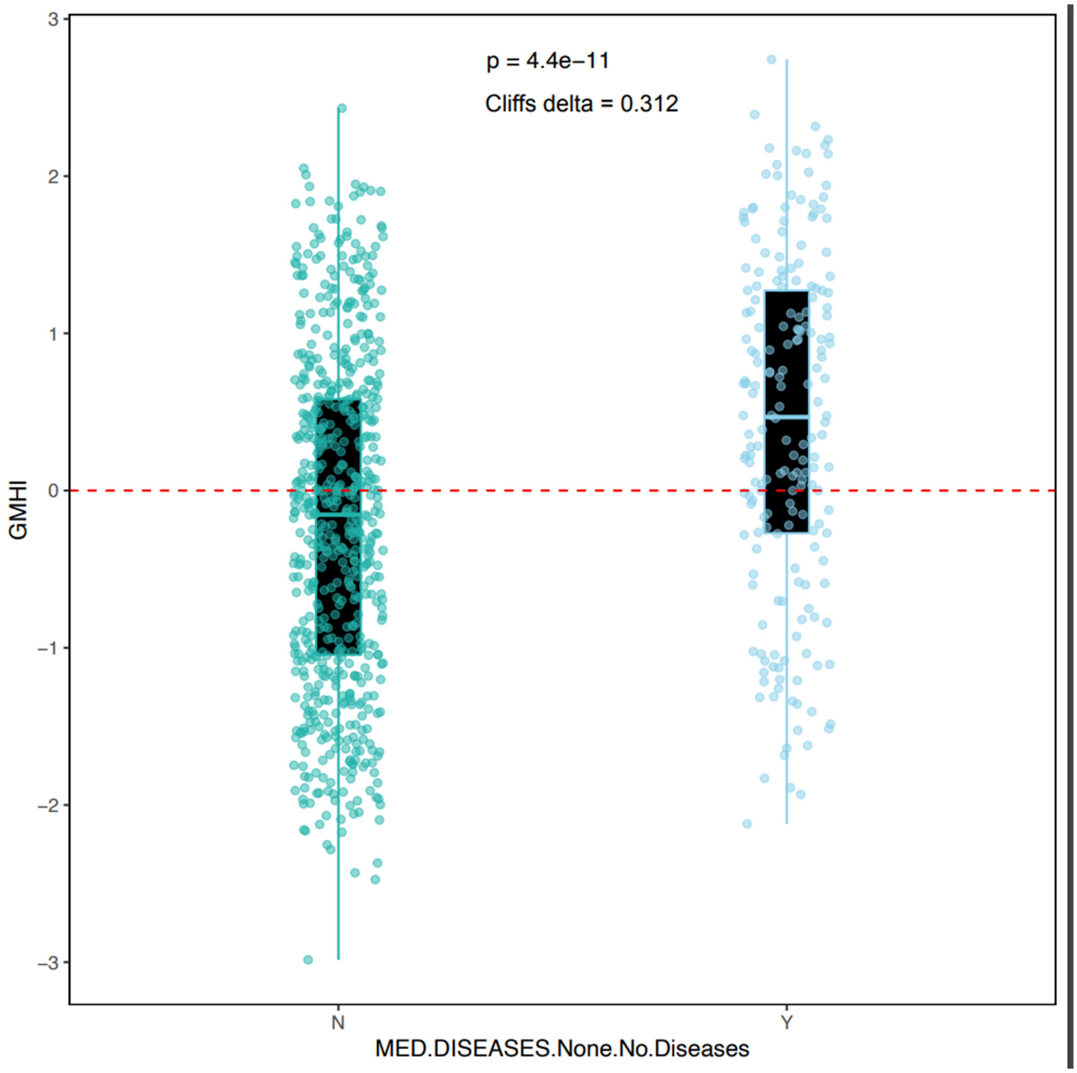


***Supplementary figure 6: Gut Microbiome Health Index calculated for DMP cohort***

*Figure shows box-plots of Gut Microbiome Health Index (GMHI) for healthy participants of DMP cohort samples (Y) vs participants who reported one or more diseases (N).*

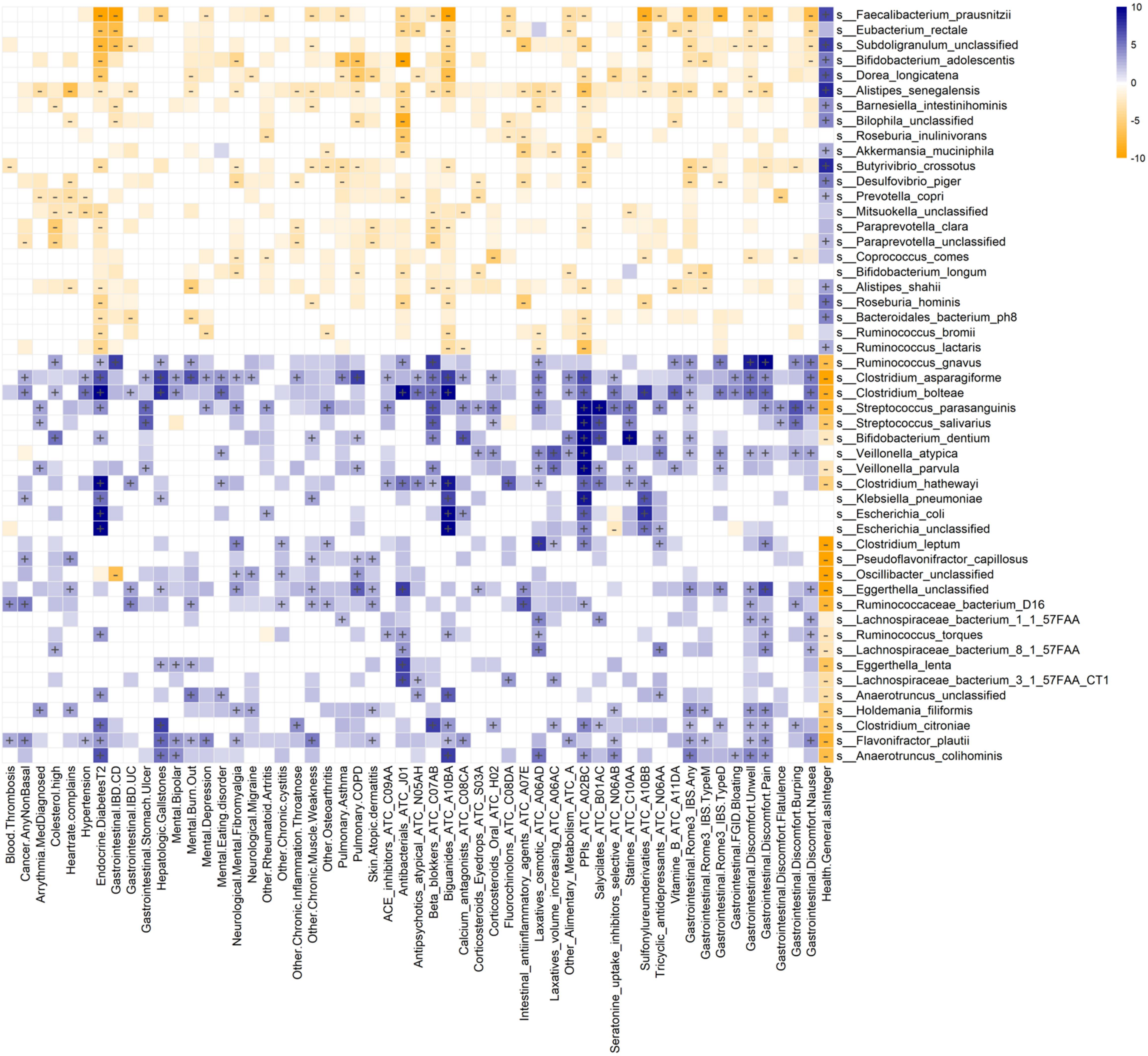


***Figure S7: Microbiome associations with diseases and medication use***

*Heatmap displays microbiome-phenotype associations, with microbial species clustered by association p-value using hierarchical clustering and colored by the direction of association. Study-wide significant associations (FDR < 0.05) are marked with +/-, while colored associations without label mark nominally significant associations (p-value < 0.05).*

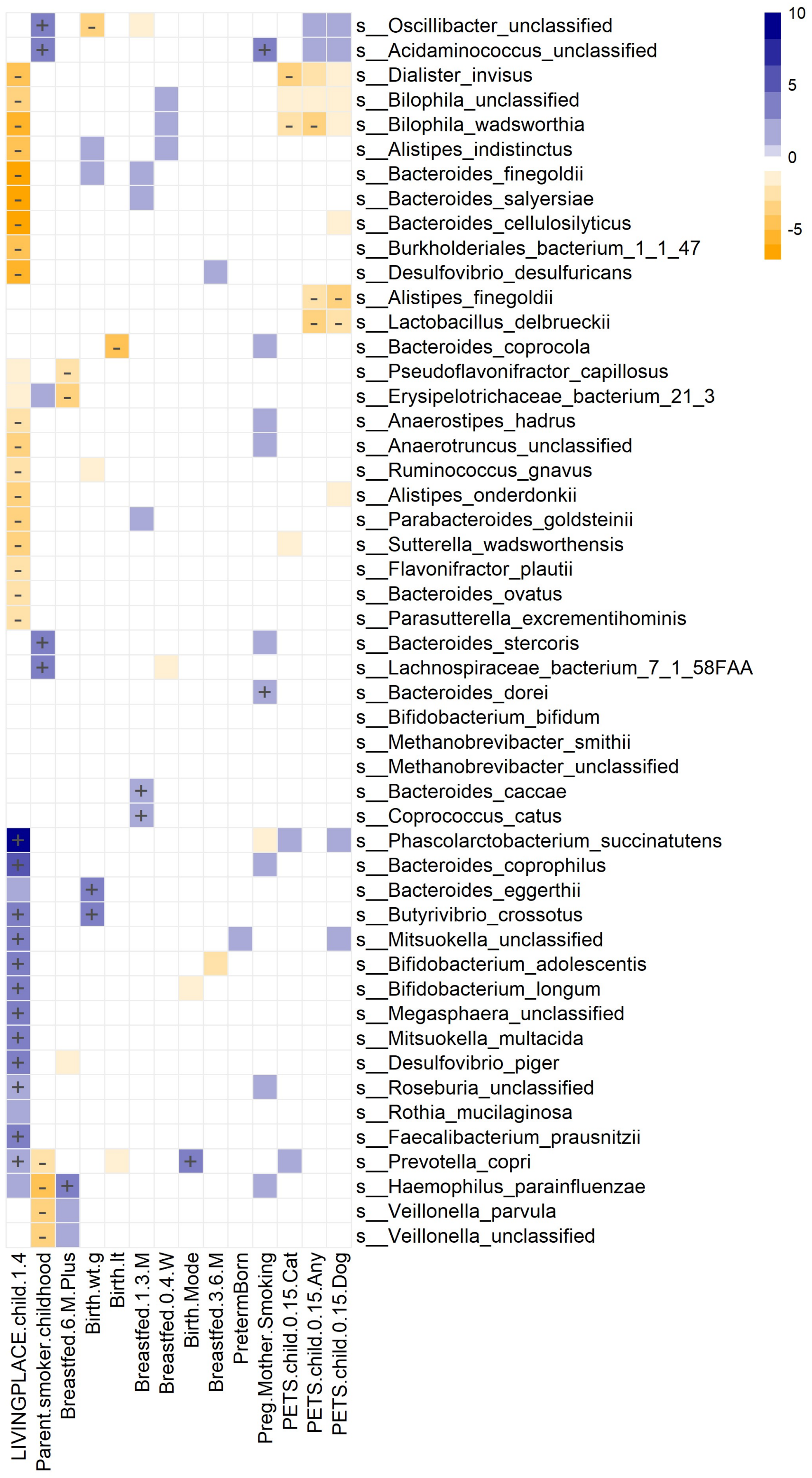


***Figure S8: Microbiome association with early-life exposures***

*Heatmap displays microbiome-phenotype associations, with microbial species clustered by association p-value using hierarchical clustering and colored by the direction of association. Study-wide significant associations (FDR < 0.05) are marked with +/-, while colored associations without label mark nominally significant associations (p-value < 0.05).*

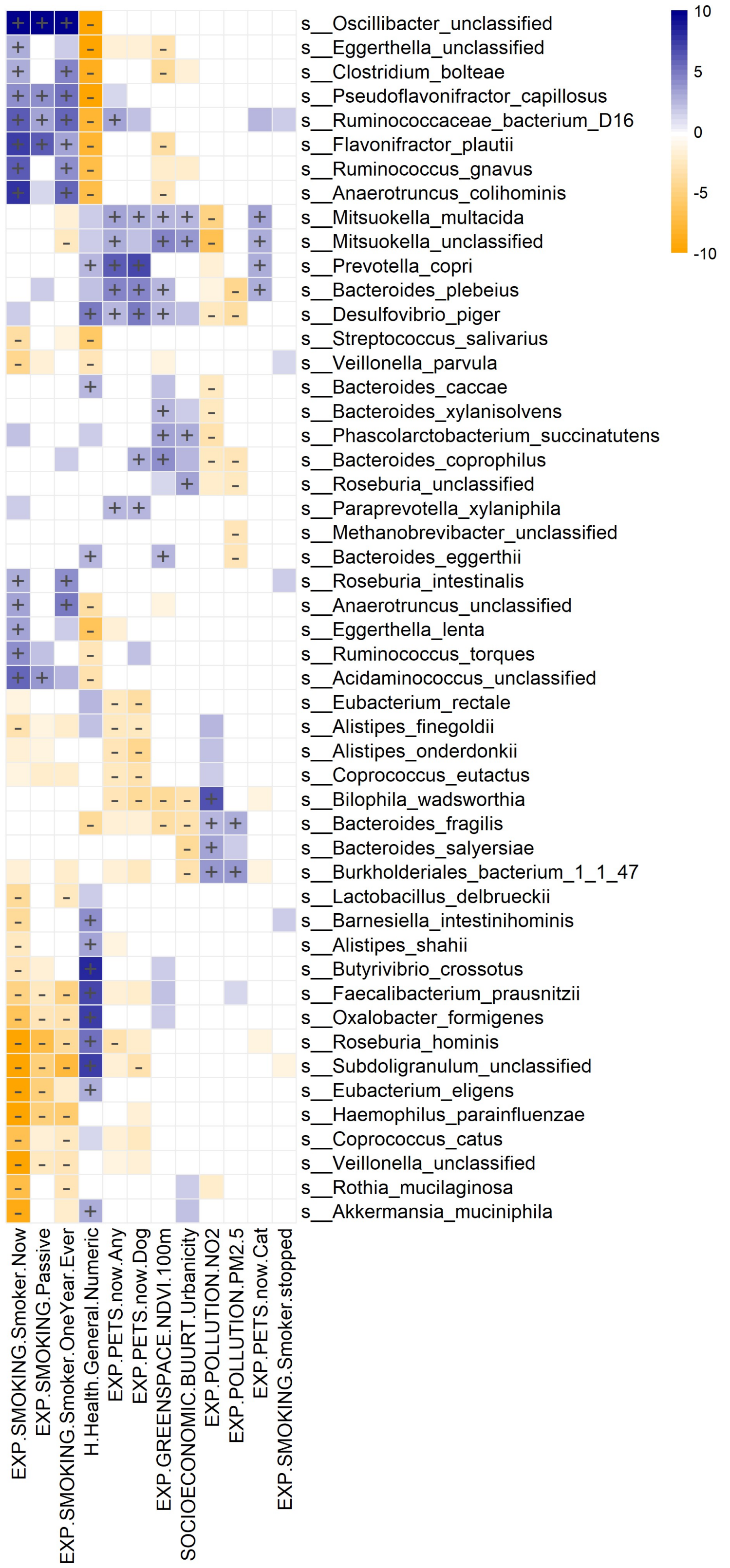


***Figure S9: Microbiome association with smoking, pollutants and greenspace***

*Heatmap displays microbiome-phenotype associations, with microbial species clustered by association p-value using hierarchical clustering and colored by the direction of association. Study-wide significant associations (FDR < 0.05) are marked with +/-, while colored associations without label mark nominally significant associations (p-value < 0.05).*

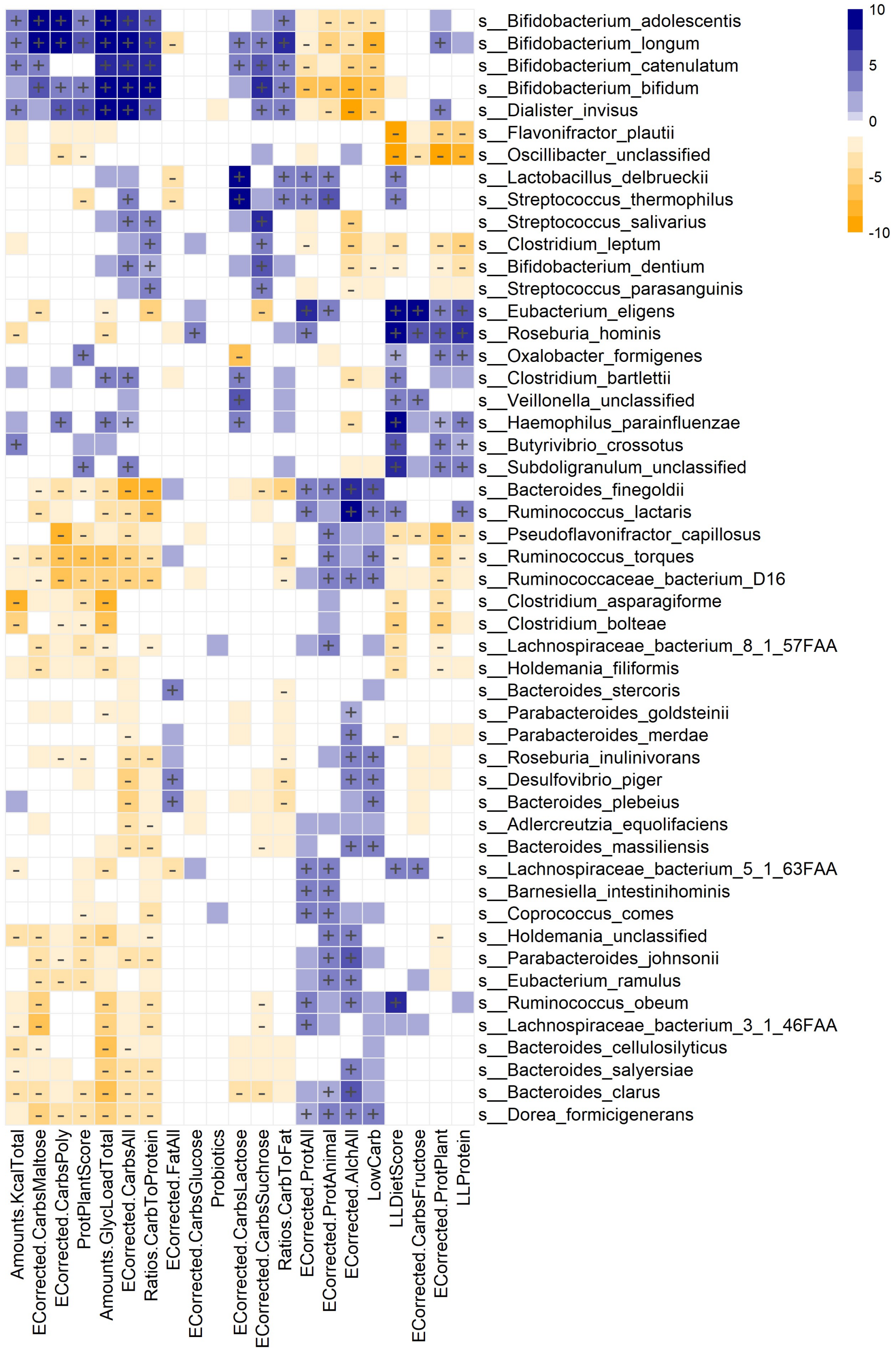


***Figure S10: Microbiome association with diet***

*Heatmap displays microbiome-phenotype associations, with microbial species clustered by association p-value using hierarchical clustering and colored by the direction of association. Study-wide significant associations (FDR < 0.05) are marked with +/-, while colored associations without label mark nominally significant associations (p-value < 0.05).*
